# Species-specific chromatin landscape determines how transposable elements shape genome evolution

**DOI:** 10.1101/2022.03.11.484033

**Authors:** Yuheng Huang, Harsh Shukla, Yuh Chwen G. Lee

## Abstract

Transposable elements (TEs) are selfish genomic parasites that increase their copy number at the expense of host fitness. The “success,” or genome-wide abundance, of TEs differs widely between species. Deciphering the causes for this large variety in TE abundance has remained a central question in evolutionary genomics. We previously proposed that species-specific TE abundance could be driven by the inadvertent consequences of host-direct epigenetic silencing of TEs—the spreading of repressive epigenetic marks from silenced TEs into adjacent sequences. Here, we compared this TE-mediated “epigenetic effect” in six species in the *Drosophila melanogaster* subgroup to dissect step-by-step the role of such effect in determining genomic TE abundance.

We found that TE-mediated spreading of repressive marks is prevalent and substantially varies across and even within species. While this TE-mediated effect alters the epigenetic states of adjacent genes, we surprisingly discovered that the transcription of neighboring genes could reciprocally impact this spreading. Importantly, our multi-species analysis provides the power and appropriate phylogenetic resolution to connect species-specific host chromatin regulation, TE-mediated epigenetic effects, the strength of natural selection against TEs, and genomic TE abundance unique to individual species. Our findings point towards the importance of host chromatin landscapes in shaping genome evolution through the epigenetic effects of a selfish genetic parasite.

## Introduction

Transposable elements (TEs) are widespread genetic parasites that copy and insert themselves across host genomes. The presence and movement of TEs could impair host genome functions. TEs disrupt genes and functional elements (Finnegan 1992), introduce ectopic regulatory sequences (Chuong *et al*. 2017), and trigger highly deleterious chromosomal rearrangements through nonhomologous recombination (Langley *et al*. 1988; Montgomery *et al*. 1991). Nevertheless, the ability to self-replicate has allowed TEs to successfully occupy nearly all eukaryotic genomes surveyed (Wells and Feschotte 2020). Intriguingly, the abundance of TEs substantially varies across the phylogenetic tree (Huang *et al*. 2012; Elliott and Gregory 2015; Wells and Feschotte 2020). For instance, the proportion of vertebrate genomes occupied by TEs ranges from only 6% in pufferfish (Volff *et al*. 2003) to more than 65% in salamander (Nowoshilow *et al*. 2018). Even within the same genus, genomic TE abundance differs widely (e.g., 2.5%-25% in *Drosophila* (Clark *et al*. 2007; Rius *et al*. 2016)). Deciphering the role of this prevalent genetic parasite in shaping genome evolution has remained a central question in genomics (Kazazian 2004; Feschotte and Pritham 2007; Arkhipova 2018); however, the ultimate causes of such dramatic divergence in TE abundance remain unclear.

Theoretical analyses proposed that, in panmictic host populations with unrestricted recombination, TE abundance is determined by how quickly TEs replicate and how fast they are removed from the populations by natural selection against their harmful fitness effects ((Charlesworth and Charlesworth 1983), reviewed in (Lee and Langley 2010)). Under this model, divergent genome-wide TE abundance could be driven by between-species differences in the strength of selection against TEs. Currently available evolutionary models that address this possibility have focused on population genetic parameters that influence the *efficacy of selection* removing TEs, such as mating systems (Wright and Schoen 1999; Dolgin and Charlesworth 2006; Boutin *et al*. 2012) and effective population size (Lynch and Conery 2003). Yet, empirical support for such hypothesis has been mixed (Dolgin *et al*. 2008; Lockton and Gaut 2010; de la Chaux *et al*. 2012; Arunkumar *et al*. 2014; Ågren *et al*. 2014; Mérel *et al*. 2021; Oggenfuss *et al*.

2021). On the other hand, between-species differences in the magnitude of harmful effects exerted by TEs, and accordingly the strength of selection against TEs, could also determine genomic TE abundance, a plausible hypothesis that is yet to have empirical investigations.

A new avenue for exploring how these genetic parasites shape the function and evolution of eukaryotic genomes is opened by the recently discovered host-directed silencing of TEs and the associated “inadvertent” deleterious epigenetic effects (reviewed in (Choi and Lee 2020)). To counteract the selfish increase of TEs in host genomes, eukaryotic hosts have evolved small RNA-mediated mechanisms to transcriptionally silence TEs (reviewed in (Slotkin and Martienssen 2007; Czech *et al*. 2018; Deniz *et al*. 2019)). Host protein complexes are guided by small RNAs to TEs with complementary sequences, which is followed by the recruitment of methyltransferases that modify DNA or histone tails at TE sequences (Qi *et al*. 2006; Aravin *et al*. 2008; Wang and Elgin 2011; Sienski *et al*. 2012; Le Thomas *et al*. 2013). Such process results in the enrichment of DNA methylation or di- and tri-methylation on lysine 9 of histone H3 (H3K9me2/3), both repressive epigenetic modifications that are typically found at heterochromatic regions and associated with repressed gene expression (reviewed in (Pikaard and Mittelsten Scheid 2014; Allis and Jenuwein 2016)). This repressed transcription of TEs results in reduced RNA intermediates (for RNA-based TEs) and proteins (e.g., transposase and reverse transcriptase) necessary for TE replication, effectively slowing the selfish propagation of TEs.

While such epigenetic silencing of TEs should benefit their hosts, studies in various model species have found that repressive marks enriched at silenced TEs “spread” to adjacent sequences across the euchromatic genomes (i.e., *Mus*, *Drosophila, Arabidopsis,* and *Oryza*; (Rebollo *et al*. 2012; Sienski *et al*. 2012; Lee 2015; Quadrana *et al*. 2016; Choi and Purugganan 2018), reviewed in (Choi and Lee 2020)).

Furthermore, TEs with such effects were observed to have lower population frequencies (Hollister and Gaut 2009; Lee 2015; Lee and Karpen 2017), suggesting that selection acts to them. These discoveries highlight the potential importance of TE-triggered epigenetic effects in shaping genome evolution. Interestingly, the strength of TE-mediated spreading of repressive marks substantially varies between distantly related taxa (reviewed in (Choi and Lee 2020)). Investigations on pairs of closely related species further revealed that this “epigenetic effect of TEs” differs and is stronger in the species with fewer TEs (*Arabidopsis thaliana v.s. A. lyrate* and *Drosophila melanogaster v.s. D. simulans*; (Hollister *et al*. 2011; Lee and Karpen 2017)). These observations spurred our previous hypothesis that varying epigenetic effects of TEs could result in between-species differences in the strength of selection against TEs, eventually contributing to divergent genomic TE abundance (Lee and Karpen 2017). We further postulated that this difference in TE-mediated epigenetic effects could have resulted from species-specific genetic modulation of the repressive chromatin landscape (Lee and Karpen 2017), which was shown to determine the spreading of repressive epigenetic marks from constitutive heterochromatin (reviewed in (Girton and Johansen 2008; Elgin and Reuter 2013)).

To fully examine the hypothesis that host chromatin landscape drives between-species differences of TEs through epigenetic mechanisms, one needs to connect species-specific regulation of chromatin landscape, TE-mediated spreading of repressive marks, the associated functional consequence and resultant selection against TEs, and genomic TE abundance. However, former analyses that compared TE-mediated epigenetic effects between species have limited sampling (two species) and thus lack sufficient statistical power for robust inference (Hollister *et al*. 2011; Lee and Karpen 2017). Also, support for key links of the hypothesis is lacking. For instance, selection against TE-mediated epigenetic effects is predicted to result from the associated reducing effects on the expression of neighboring genes. Yet, investigations in multiple taxa reported weak or no associations between the epigenetic effects of TEs and neighboring gene expression ((Quadrana *et al*. 2016; Stuart *et al*. 2016; Lee and Karpen 2017), reviewed in (Kelleher *et al*. 2020; Choi and Lee 2020)). These inconclusive observations cast doubt on the possibility that TE-mediated epigenetic effects impair host fitness by silencing neighboring genes and whether this particular deleterious consequence indeed shapes genome evolution. Furthermore, previous comparisons of population frequencies between TEs with and without epigenetic effects, an approach used to infer the strength of natural selection removing TEs, could not exclude the confounding influence of other harmful effects of TEs on their population frequencies (e.g., (Hollister and Gaut 2009; Lee 2015)). Accordingly, those analyses could not unequivocally support selection against TE-mediated epigenetic effects. Multi-species studies that span an appropriate evolutionary distance and connect the missing links in the proposed hypothesis would be needed to test the predicted importance of TE-mediated epigenetic effects in determining between-species differences of TEs.

In this study, we investigated the prevalence of TE-mediated local enrichment of repressive epigenetic mark, H3K9me2, in the euchromatic genome of six species in the *Drosophila melanogaster* subgroup (diverged around 10 MYR, (Obbard *et al*. 2012), **Figure 1A**). These species are from the two well-studied species complexes (*melanogaster* and *yakuba* complexes), providing good phylogenetic resolution to address the role of TE-mediated epigenetic effects in genome evolution. While TE insertions in all species studied result in robust local enrichment of repressive epigenetic marks, the strength of such effects varies substantially within genomes, between species, and among species complexes. Our larger sample size allowed us to re-examine the still debated question about the impacts of TE-mediated enrichment of repressive marks on neighboring gene expression, which surprisingly revealed their complex interactions. Importantly, our multi-species analysis provides the power to test the predicted associations between genomic TE abundance, TE-mediated epigenetic effects, selection against such effects, and host chromatin environment, while uncovering the evolutionary causes for the wide variety of TE profiles between species.

**Figure 1.**
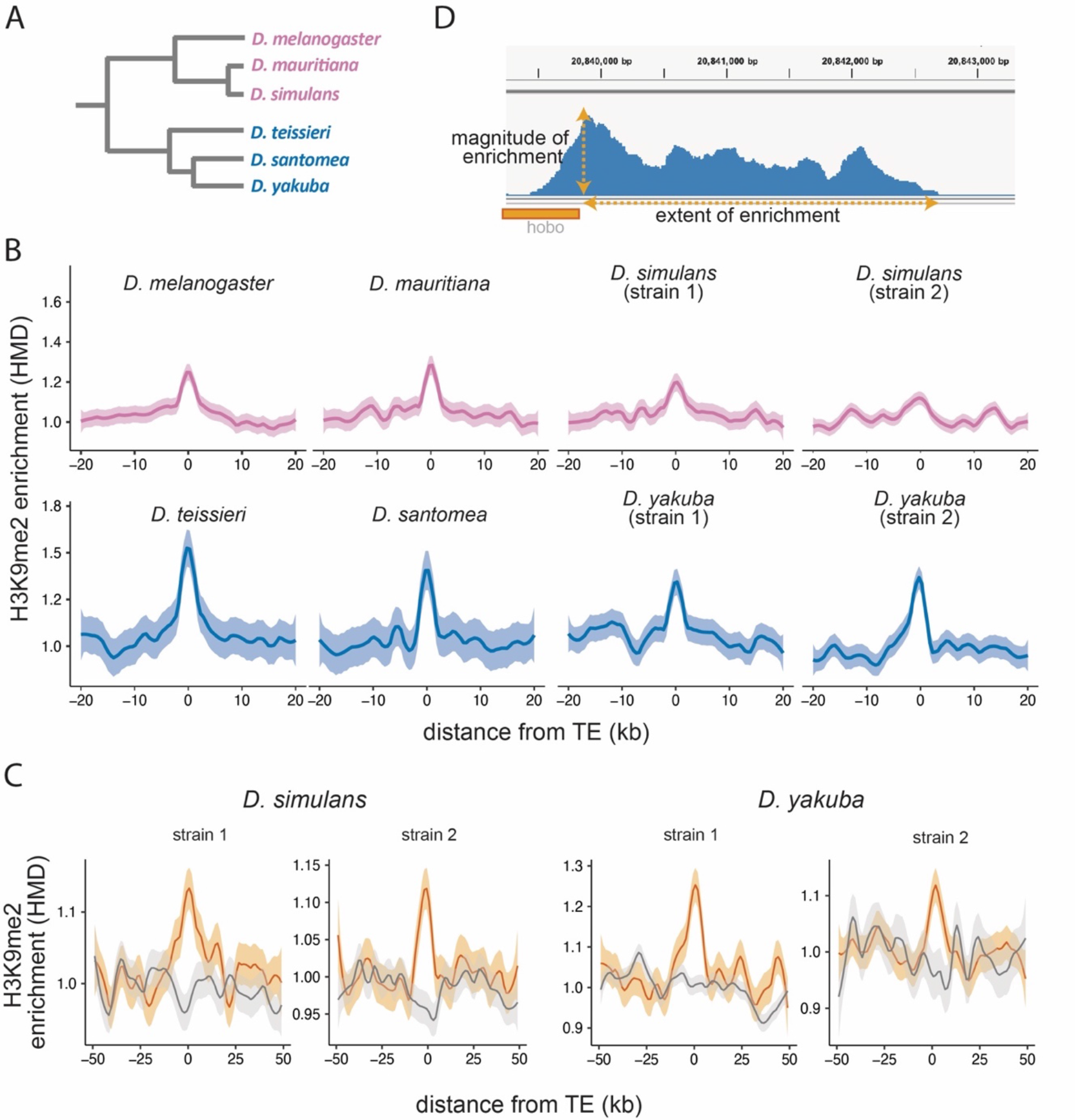
Enrichment of H3K9me2 around euchromatic TEs across species. **(A)** Phylogenetic relationship among species included in this study. Species in the *D. melanogaster* complex are in pink while those in *D. yakuba* complex are in blue. **(B)** Genome-wide average H3K9me2 HMD levels around euchromatic TEs with LOESS smoothing (span = 15%) in studied genomes. 95% confidence interval around smooth is shown in shaded area. **(C)** Genome-wide average H3K9me2 HMD levels at homologous sequences in the presence (orange) and absence (gray) of euchromatic TEs in species that have data for two strains (*D. simulans* an*d D. yakuba*). The average of H3K9me2 HMD level was smoothed with LOESS (span = 15%) with 95% confidence intervals around smooth shown in shaded area. (D) An Integrated-Genome-Viewer view showing the local enrichment of H3K9me2 around a *hobo* TE, and the two estimates (magnitude and extent of H3K9me2 enrichment) estimated for quantifying the epigenetic effects of individual TE.

## Results

### TE-mediated *cis*-spreading of repressive marks is prevalent across studied species

We investigated whether previously reported local enrichment of heterochromatic marks around euchromatic TEs in model *Drosophila* species (Lee 2015; Lee and Karpen 2017) is prevalent across species in the *Drosophila melanogaster* subgroup. To do so, we performed spike-in controlled Chromatin immunoprecipitation (ChIP)-seq targeting H3K9me2, a histone modification that is highly enriched in the constitutive heterochromatin in *D. melanogaster* (Riddle *et al*. 2011; Kharchenko *et al*. 2011) (see Materials and Methods). We estimated histone modification density (HMD), which is the ratio of fragment coverage in ChIP samples to that in matching input, standardized by the ratio of spike-in fragments (Lam *et al*. 2019) (see Materials and Methods). TE insertions in strains used for the ChIP-seq experiment were annotated by running Repeatmodeler (Flynn *et al*. 2020) on PacBio genomes, whose high continuity enables more comprehensive identification of TEs than previous studies based on short-read sequencing data (Khost *et al*. 2017; Chakraborty *et al*. 2019). Because heterochromatic regions of the genome are already enriched with H3K9me2, we excluded TEs in or near heterochromatic regions from our analysis (see Materials and Methods).

Across all six species analyzed, we observed significant enrichment of H3K9me2 HMD around TEs, and this enrichment decreases to the background level within 10kb (**Figure 1B**). To exclude the possibility that this enrichment of H3K9me2 is due to TE preferentially inserted into regions that are already enriched with heterochromatic marks (Dimitri and Junakovic 1999), we collected H3K9me2 epigenomic data for two genomes of *D. simulans and D. yakuba* and compared the enrichment of H3K9me2 for homologous sequences with and without a TE insertion. The H3K9me2 enrichment is observed in the vicinity of TEs in the genome where they are present, but not at homologous sequences in the other genome (**Figure 1C**). This observation expanded previous single-species observations (Lee 2015; Lee and Karpen 2017) and supports that the local enrichment of heterochromatic marks in the euchromatic regions are induced by TEs in multiple species.

### Strength of TE-mediated epigenetic effects depends jointly on TE attributes and host genetic background

In order to investigate biological factors associated with the strength of TE-mediated epigenetic effects, we quantified the local enrichment of H3K9me2 for individual TEs. Specifically, we estimated two indexes, the magnitude and the extent of H3K9me2 enrichment (**Figure 1D**, see Materials and Methods), and found them highly correlated within individual genomes (*Spearman rank correlation coefficients (⍴)* = 0.52-0.69, *p <* 10^-16^, **Figure 1 – Supplementary Figure S1**). We also estimated the magnitude and extent of TE-induced H3K9me2 enrichment by comparing two genomes of the same species (*D. simulans* and *D. yakuba*; see Materials and Methods) and found that estimates based on one genome or two genomes strongly correlate (*Spearman rank correlation coefficients (⍴)* = 0.64-0.85 (magnitude) and 0.40-0.64 (extent), *p <* 10^-10^ for all tests, **Figure 1 – Supplementary Figure S2**). Because an important aspect of our analyses is the comparison of TE-induced H3K9me2 across species, we reported analyses based on single genome henceforth.

TE length was postulated to be an important factor determining the strength of TE-mediated epigenetic effects because silenced TEs that are longer in length are expected to represent larger heterochromatin mass (Lee 2015). Consistent with the prediction, we observed significant, though weak, positive correlations between TE length and the strength of TEs’ epigenetic effects within most genomes studied (*Spearman rank correlation coefficients (⍴)* = 0.12-0.22, *p <* 0.05; **Figure 1–Supplementary Figure S3**). It is worth noting that previous analysis on non-reference *D. melanogaster* strains was unable to test this prediction, due to the inability to assemble internal sequences of TEs with short-read resequencing data (Lee and Karpen 2017).

We next compared TEs of different classes, which are classifications based on the transposition mechanisms of TEs (Wicker *et al*. 2007) and previously observed to associate with varying extent of epigenetic effects within *D. melanogaster* genomes (Lee 2015; Lee and Karpen 2017). While different classes of TEs showed similar levels of H3K9me2 enrichment, there is a very strong species-complex effect in which TEs in genomes of *yakuba* complex showed much larger *magnitude* of epigenetic effects than those in genomes of *melanogaster* complex (**Figure 2A**, an average 1.7-fold larger). On the other hand, the *extent* of H3K9me2 enrichment is more variable and does not show a similar trend (**Figure 2A**). Interestingly, the extent and magnitude of H3K9me2 spreading varies substantially between TEs of different families within a class and between TEs of the same family (**Figure 2B** for *D. simulans* and see **Figure 2 – Supplementary Figure S1** for other *melanogaster* complex species and **Figure 2 – Supplementary Figure S2** for *yakuba* complex species). Moreover, the rank order of the extent and magnitude of TE-induced H3K9me2 enrichment of TE families varies between species and even between strains of the same species. These observations strongly suggest that the strength of TE-mediated epigenetic effects depends on both TE family attributes and host genetic background. It is worth noting that the percentage of TEs with a family assigned is higher in the *melanogaster* complex than that in the *yakuba* complex (**Figure 2B – Supplementary Figure S3**), and our analysis likely missed TE families that are highly divergent in and/or unique to the *yakuba* complex species.

**Figure 2.**
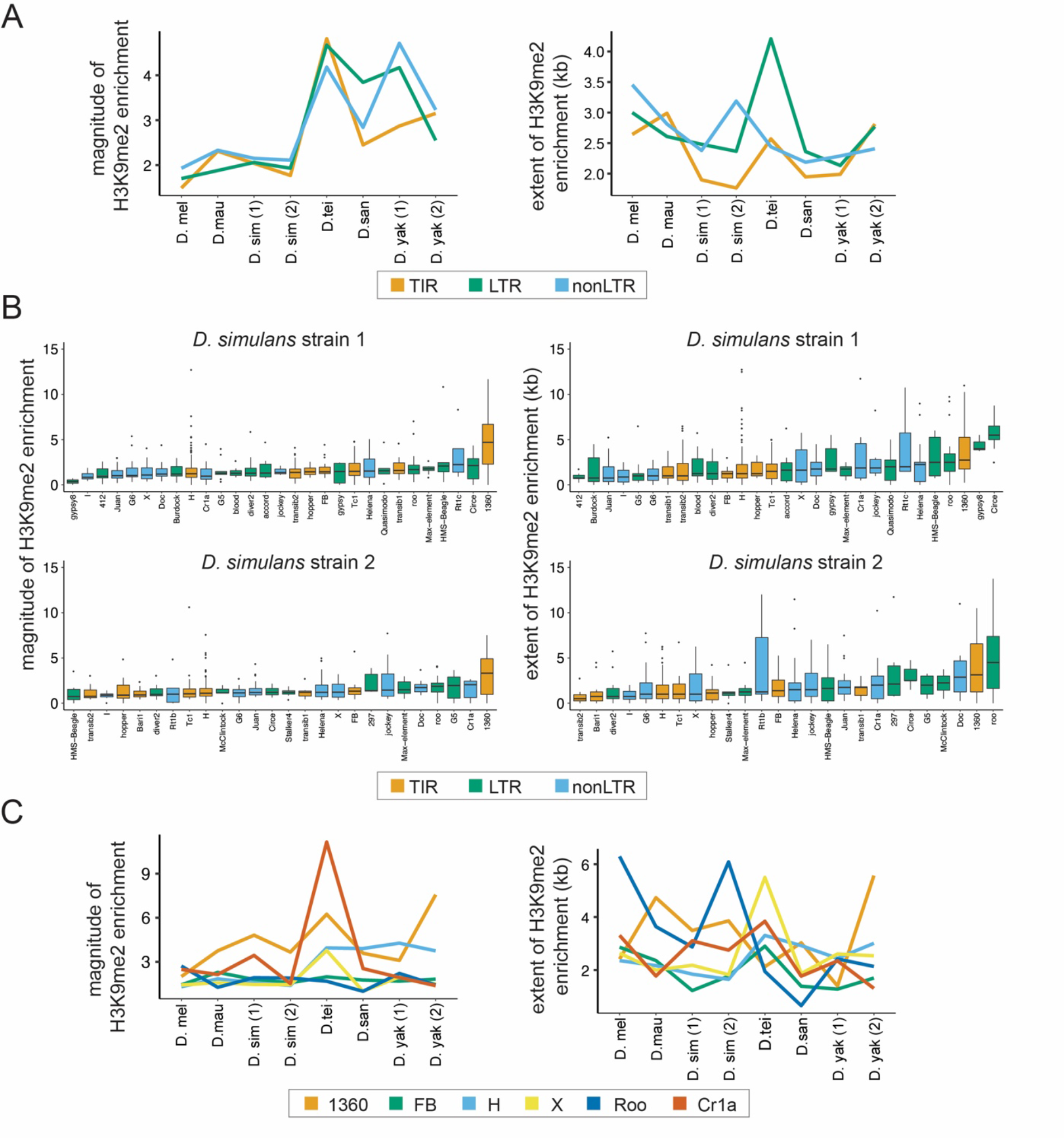
Variation in the epigenetic effects of TEs within genomes. **(A)** The mean magnitude (left) and extent (right) of TE-induced H3K9me2 enrichment for different types of TEs in eight genomes from six species are shown. Different colors represent TEs of different classes, including Terminal Inverted Repeat (TIR), Long Terminal Repeat (LTR), and non-Long Terminal Repeat (non-LTR, also known as LINE) insertions. **(B)** The magnitude (left) and extent (right) of TE-induced H3K9me2 enrichment for different TE families in the two strains of *D. simulans*. Only TE families with at least five identified copies in a genome were included. See Figure 2 **– Supplementary Figure 2 and 3** for other genomes. **(C)** The median magnitude (left) and extent (right) of TE-induced H3K9me2 enrichment for six TE families that have at least five copies in all genomes studied.

To further study the effects of host genome-by-family interaction across species, we compared the epigenetic effects of TEs from families that are shared across all strains and have at least five copies (**Figure 2C**). Using a linear regression model, we found significant family-by-strain interaction effects for both the magnitude (*ANOVA F-value: 3.4, df = 35, p = 1.4x10^-10^)* and the extent *(ANOVA F-value: 4.4, df = 35, p < 5.8x10^-16^*) of H3K9me2 enrichment. While the significant host genome-by-family interaction on the *magnitude* of H3K9me2 enrichment is likely mainly driven by Cr1a family, all TE families investigated showed strong host genome-by-family interactions for the *extent* of H3K9me2 enrichment (**Figure 2C**). Together, our observations revealed that the epigenetic effects of TEs *jointly* depend on the class, family identity, and length of TEs as well as host genetic background (also see below).

### Complex relationship between TE-induced enrichment of H3K9me2 and neighboring gene expression

Euchromatic TEs are interspersed with actively transcribing genes, and TE-mediated spreading of H3K9me2 was previously observed to extend into neighboring genes (Lee 2015; Lee and Karpen 2017). The enrichment of H3K9me2, a repressive histone modification, is generally associated with suppressed gene expression (Kouzarides 2007). Accordingly, TE-mediated epigenetic effects were expected to lower neighboring gene expression. Yet, previous studies in several model species found contradicting effects of TE-induced enrichment of repressive marks on adjacent gene expression (reviewed in (Kelleher *et al*. 2020; Choi and Lee 2020)). These observations left an important yet unsolved question about the functional importance of TEs’ epigenetic effects.

To revisit this question with expanded data, we first studied the associations between TE-mediated epigenetic effects and the *epigenetic states* of TE-adjacent genes. Similar to previous observations made in *D. melanogaster* (Lee 2015; Lee and Karpen 2017), H3K9me2 enrichment level at gene body is positively correlated with the magnitude and extent of TE-induced H3K9me2 enrichment in all species analyzed (*Spearman rank correlation coefficient* (*⍴*) = 0.11-0.32, *p <* 0.05 for all except one test, **Figure 3 – Supplementary Figure S1**). Importantly, for the magnitude of TE-mediated epigenetic effects, this association is much stronger for genes close to TEs than those far from TEs (likelihood ratio tests for regression analysis, *p* < 0.05 for all genomes except for *D. mauritiana* (*p =* 0.07) and *D. teissieri* (*p =* 0.14); **Figure 3A**), which echoes the observation that TE-mediated H3K9me2 enrichment is distance-dependent (**Figure 1B**). For the extent of TE-mediated epigenetic effects, we observed similar, but less consistent, distance-dependent effects (likelihood ratio tests for regression analysis*, p* < 0.05 for all genomes except for *D. simulans* (strain 2, *p* = 0.97) and *D. santomea* (*p* = 0.11); **Figure 3 – Supplementary Figure S2**). Such observation could be because, instead of quantifying the magnitude of H3K9me2 enrichment, the extent index measures other aspects of TE-mediated epigenetic effects (also see below). On the contrary, we found no significant correlations between TE-mediated H3K9me2 enrichment and the *expression rank* of adjacent genes (*Spearman rank correlation tests, p >* 0.05 for all genomes). Furthermore, the associations between gene expression and the epigenetic effects of TEs do not differ between close and distant gene-TE pairs (likelihood ratio tests for regression analysis, *p* > 0.05 for all genomes, **Figure 3B** and **Figure 3 – Supplementary Figure S3** for the magnitude and extent of TE-mediated epigenetic effects, respectively), which is contrary to what we observed for TE-mediated effects on genic epigenetic states (**Figure 3A**).

**Figure 3.**
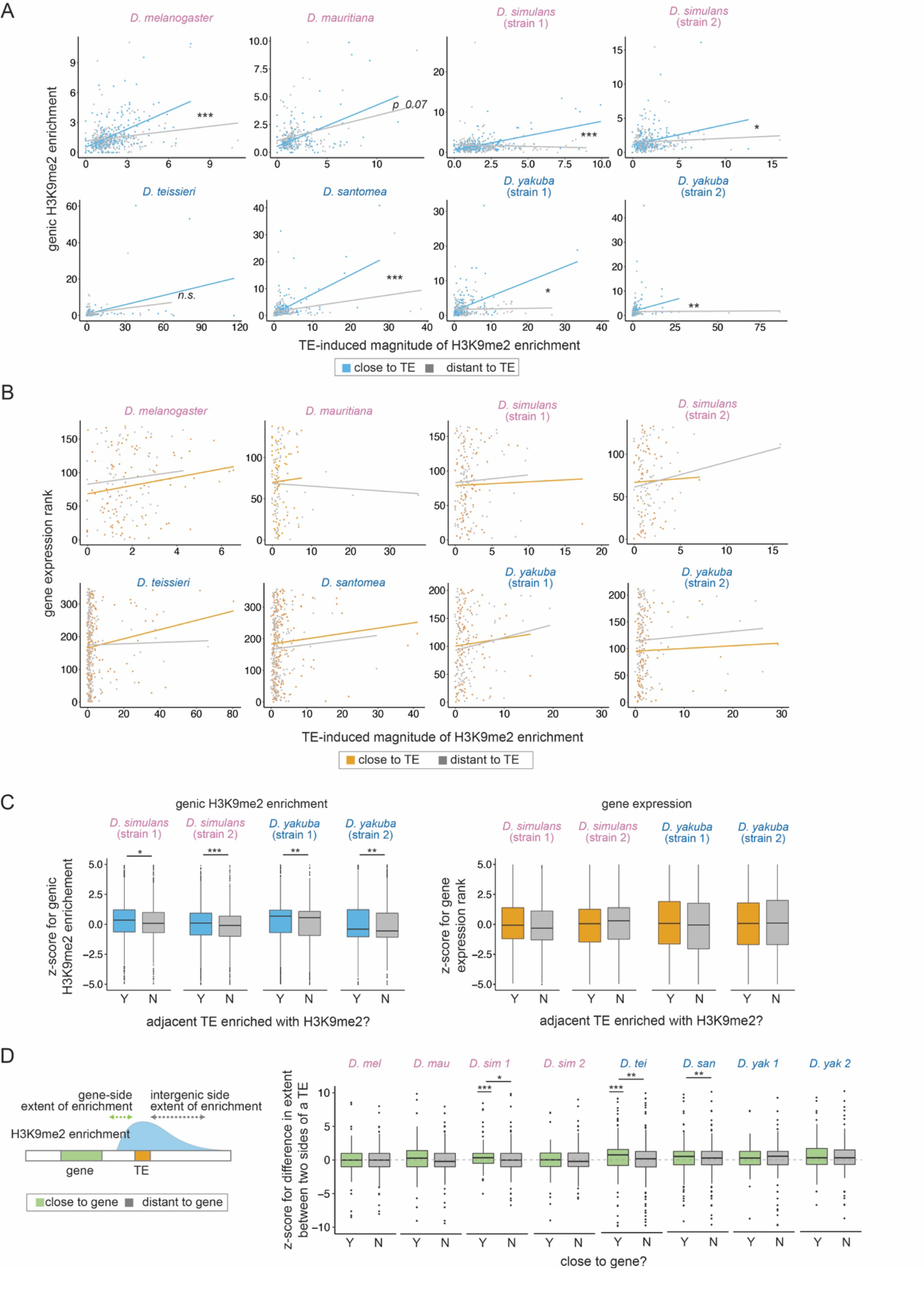
Associations between TE-mediated H3K9me2 enrichment and the epigenetic states and expression of neighboring genes. **(A)** The associations between the magnitude of TE-mediated H3K9me2 enrichment and genic H3K9me2 enrichment are much stronger for genes close to TEs (distance to a TE is smaller than the 50% quantile, blue) than for genes distant to TEs (gray; see Figure 3 **– Supplementary** Figure 2 for the extent of TE-mediated H3K9me2 enrichment). **(B)** The associations between the magnitude of TE-mediated H3K9me2 enrichment and gene expression rank *does not* differ between genes close (orange) and distant (gray) to TEs (see Figure 3 **– Supplementary** Figure 3 for the extent of TE-mediated H3K9me2 enrichment). **(C)** Z-scores for comparing the H3K9me2 enrichment (left) and expression rank (right) of homologous genic alleles whose nearby TEs with (blue/orange) or without (gray) epigenetic effects (as defined as the magnitude of H3K9me2 enrichment > 1; see Figure 3 **– Supplementary** Figure 4 for categorizing TEs with the extent of H3K9me2 enrichment, which gives similar results). Positive Z-score means the allele with TE has higher H3K9me2 enrichment or larger expression rank (i.e., lower expression level) than the homologous allele without TE in another strain. **(D)** A cartoon describing the “gene side” and “intergenic side” extent of H3K9me2 enrichment mediated by TEs (left). Z-scores for comparing the extent of TE-mediated H3K9me2 enrichment on the intergenic side and on the genic side for TEs close (green) and far (gray) from genes whose expression is at least 10 RPKM. Positive z-score means that the extent of spread is more restricted on the genic side than the intergenic side. In several genomes, z-scores for TEs close to genes are significantly different from zero and/or larger than those for TEs distant to genes. \*\*\**p <* 0.001, \*\**p <* 0.01, \**p<* 0.05 for *likelihood ratio tests* (A, B) and *Mann-Whitney tests* (C, D).

The above analyses may have limited power because variation in gene expression levels within a genome could be due to intrinsic gene properties, instead of TE-mediated effects. To directly test the effects of TE-mediated H3K9me2 enrichment on nearby gene expression, we compared the expression of homologous alleles with and without adjacent TEs in *D. simulans* and *D. yakuba,* two species that we have epigenomic and transcriptomic data for two strains. We estimated z-scores, which compare H3K9me2 enrichment and expression rank between homologous alleles (see Materials and Methods). A positive z-score indicates that the allele adjacent to TE insertions has higher H3K9me2 enrichment or expression rank (i.e., lower expression) than the homologous allele without a nearby TE. We again observed TEs with stronger epigenetic effects associate with higher z-score of genic H3K9me2 enrichment, which confirms their impacts on genic epigenetic states (*Mann-Whitney U test, p <* 0.05 for all comparisons; **Figure 3C** and **Figure 3 – Supplementary Figure S4** for categorizing genes according to the magnitude and extent of TE-induced H3K9me2 enrichment, respectively). On the contrary, there is no association between TE-mediated epigenetic effects and z-score of gene expression rank (*Mann-Whitney U test, p >* 0.05 for all comparisons; **Figure 3C** and **Figure 3 – Supplementary Figure S4** for categorizing genes according to the magnitude and extent of TE-induced H3K9me2 enrichment, respectively). Overall, our results suggest that TE-mediated epigenetic effects lead to robust enrichment of heterochromatic marks at neighboring genes, which, contrary to expectation, does not reduce the expression of neighboring genes.

The extent of local enrichment of repressive epigenetic marks for a handful of TE insertions was previously suggested to be influenced by the expression of neighboring genes in a mouse cell line (Rebollo *et al*. 2012). This observation suggests the possibility that the extent of TEs’ epigenetic effects is, in return, restrained by the expression of neighboring genes. If true, this phenomenon may explain our observed lack of negative associations between TE-mediated epigenetic effects and neighboring gene expression. To investigate this possibility on a genome-wide scale, we compared the H3K9me2 enrichment for a TE on the side facing a gene with appreciable expression in 16-18hr embryo (at least 10 RPKM; gene side) and the other side that does not face the gene (intergenic side; **Figure 3D**). We predict that the transcriptional effects of genes on TE-mediated enrichment of H3K9me2 should be the most prominent on the “gene side” of a TE. Also, the differences between the two sides of a TE should be larger for TEs closer to genes.

To test these predictions, we estimated the normalized difference between TE-mediated H3K9me2 enrichment on the intergenic and genic side by calculating a z-score. A positive z-score would mean that the extent or magnitude of TE-mediated H3K9me2 enrichment is more restricted on the genic side than the intergenic sides. Consistent with our predictions, there is a general trend that the z-score for the extent of spread is positive for TEs close to highly expressed genes (RPKM > 10), although the comparisons are statistically significant only for a subset of the genomes (*Mann-Whitney test, p <* 0.001 for *D. simulans* strain 1 and *D. teissieri,* **Figure 3D**). On the other hand, z-scores for the extent of spread are not significantly different from zero for TEs far from highly expressed genes (*Mann-Whitney test, p >* 0.05 for all genomes, **Figure 3D**) and are significantly larger for TEs close to highly expressed genes than for those far from highly expressed genes (*Mann-Whitney test, p <* 0.05 for *D. simulans* strain 1, *D. teissieri,* and *D. santomea;* **Figure 3D**). Importantly, z-scores for TEs close to lowly expressed genes (RPKM < 10) are *not* significantly different from zero nor differ between TEs close or far from genes (*Mann-Whitney test, p >* 0.05 for all genomes, **Figure 3 – Supplementary Figure S5**). Curiously, comparisons based on the *magnitude* of TE-induced H3K9me2 enrichment did not find differences between genic side and the intergenic side, nor between TEs that are close or far from highly expressed genes (*Mann-Whitney test*, *p >* 0.05 for all comparisons, **Figure 3 – Supplementary Figure S6**, also see Discussions). These observations reveal that the extent, but not the magnitude, of TE-mediated H3K9me2 spreading is restricted on the TE side facing a nearby, highly expressed gene.

Even though we found no evidence for the associations between TE-mediated epigenetic effects and adjacent gene expression (**Figure 3B and 3C**), it is plausible that, for some genes, TE-induced H3K9me2 enrichment perturbed their expression. Accordingly, to exclude the potential confounding effects of TEs on the categorization of genes according to expression, it would be helpful to classify genes as highly or lowly expressed (RPKM >=10) using the transcriptome of another strain with different TE insertions. To do so, we used modEncode developmental transcriptome data (Graveley *et al*. 2011), which were generated from a *D. melanogaster* strain that has different TE insertions (annotated in (Rahman *et al*. 2015)) from our focused strain. We categorize genes into those with high (RPKM >=10) and low (RPKM < 10) expression at 16-18 hr embryos, the same developmental stage as our epigenomic and transcriptomic data (see Materials and Methods). Similar to our other analyses, we found significantly shorter extent of H3K9me2 spread on the gene side than the intergenic side for TEs near highly expressed genes (*Paired Mann-Whitney test*, *p <* 0.05), but not for TEs near lowly expressed genes (*Mann-Whitney test*, *p >* 0.05, **Figure 3 – Supplementary Figure S7)**. Overall, our findings reveal a complex relationship between TE-mediated epigenetic effects and neighboring gene expression, and strongly suggest that the extent of TE-mediated H3K9me2 spreading is influenced by the expression of adjacent genes, especially the side of a TE facing genes.

### TEs with epigenetic effects are selected against across species

TEs exerting epigenetic effects were previously suggested to experience stronger purifying selection than other TEs (reviewed in (Choi and Lee 2020)). If TE-mediated epigenetic effects indeed impair host fitness and are thus selected against, such effects could play important roles in shaping the evolution of both TEs and their host genomes. Yet, previous analyses testing the presence of selection against TEs with epigenetic effects were restricted to few model species. More importantly, the confounding effects of other deleterious mechanisms of TEs (e.g., ectopic recombination; see below) on TE population frequencies could not be ruled out in these studies (Hollister and Gaut 2009; Lee 2015; Lee and Karpen 2017).

To investigate the fitness impacts of TE-mediated H3K9me2 enrichment, we estimated the frequencies of TEs that are included in our analysis in four species, using previously published populations of genomes (*D. melanogaster* (Lack *et al*. 2015)*, D. simulans and D. yakuba* (Rogers *et al*. 2014), and *D. mauritiana* (Garrigan *et al*. 2014)). Population frequencies of TEs are strongly influenced by the strength of natural selection against their deleterious effects (reviewed in (Charlesworth and Langley 1989; Lee and Langley 2010; Barrón *et al*. 2014)), with low-frequency TEs generally expected to be more deleterious than high-frequency TEs. We found that the *magnitude* of TE-induced H3K9me2 enrichment is significantly higher for low-frequency TEs than for high-frequency TEs in all four species (*Mann-Whitney U test, p <* 0.05 for all but *D. simulans* strain 1, **Figure 4A**), which extends previous investigations in *D. melanogaster*. Intriguingly, the difference is much weaker when comparing the *extent* of TE-mediated H3K9me2 enrichment, and the comparison is significant for only one genome (*Mann-Whitney U test, p <* 0.05 for strain 2 of *D. simulans,* but > 0.05 for all other genomes; **Figure 4B**, see Discussions).

**Figure 4.**
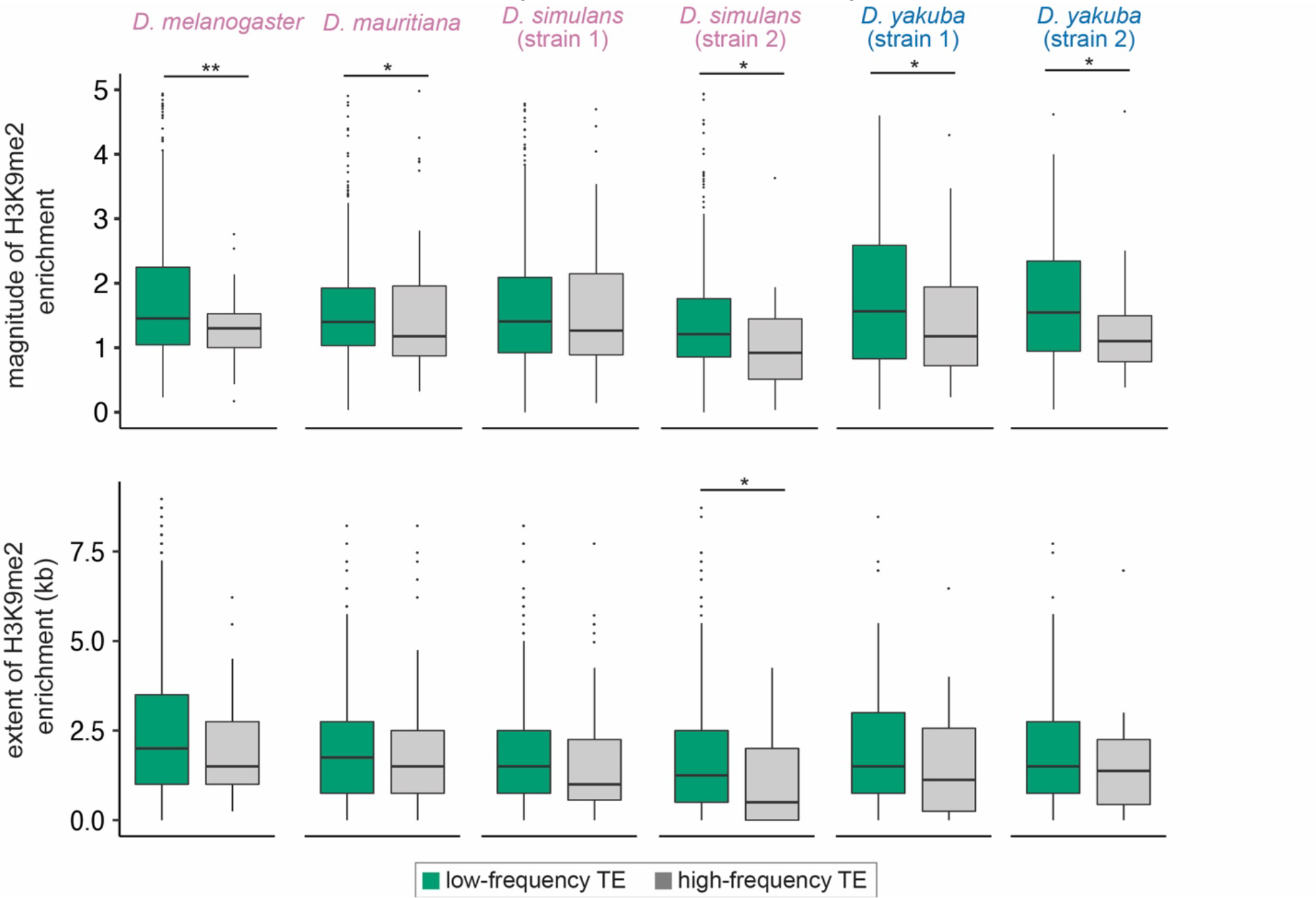
Associations between the population frequencies and epigenetic effects of TEs. The magnitude (Top) and extent (Bottom) of TE-mediated enrichment of H3K9me2 for TEs that are rare in populations (usually considered as strongly selected, green) and common TEs with high frequency in populations (gray) in *D. melanogaster, D. mauritiana, D. simulans,* and *D. yakuba*. *Mann-Whitney U test,* \*\**p <* 0.01, \**p* < 0.05.

A potential confounding factor for our observed associations between TEs’ epigenetic effects and population frequencies is TE length. TE length was previously observed to negatively correlate with population frequencies of TEs (Petrov *et al*. 2003, 2011), either because of the larger potential of long TEs in disrupting functional elements or their higher propensity to be involved in deleterious ectopic recombination (Petrov *et al*. 2003). Because the strength of TE-mediated epigenetic effects also positively correlated with TE length (**Figure 1 – Supplementary Figure S3**), our observed negative associations between TE frequencies and epigenetic effects could instead result from other harmful effects of TEs. To investigate this possibility, we performed logistic regression analysis to test the effects of TE-mediated H3K9me2 enrichment on population frequencies while accounting for the influence of TE length (population frequency ∼ length + epigenetic effects). Because of the co-linearity between predictor variables in the regression model (TE length and epigenetic effects, **Figure 1 – Supplementary Figure S3**), this analysis is expected to have restricted statistical power. Nevertheless, regression coefficients for the magnitude of H3K9me2 enrichment are negative for all but one genome (**Table 1)** and the coefficient is significantly negative for *D. melanogaster*.

**Table 1.** Generalized linear regression coefficients for the effects of TE size and H3K9me2 enrichment (magnitude and extent) on the population frequencies of TEs. Negative regression coefficients for TEs’ epigenetic effects on TE population frequencies are in bold. *p < 0.05.

|  | magnitude of H3K9me2 enrichment |  | extent of H3K9me2 enrichment |  |
| --- | --- | --- | --- | --- |
|  | TE size | magnitude | TE size | extent |
| *D. melanogaster* | -4.95E-04 | **-4.59E-01\*** | -5.22E-04 | **-8.63E-05** |
| *D. mauritiana* | -1.83E-03 | **-2.74E-02** | -1.87E-03 | 1.55E-05 |
| *D. simulans* (strain 1) | -2.56E-04 | 1.07E-01 | -2.52E-04 | 9.00E-06 |
| *D. simulans* (strain 2) | -3.13E-03 | **-2.50E-01** | -3.21E-03 | **-2.41E-04** |
| *D. yakuba* (strain 1) | -3.63E-04 | **-1.68E-01** | -3.73E-04 | **-7.77E-05** |
| *D. yakuba* (strain 2) | -4.66E-04 | **-1.27E-01** | -4.72E-04 | **-2.87E-04** |

The average deleterious effects of TE insertions were found to differ between TE families (reviewed in (Charlesworth and Langley 1989; Barrón *et al*. 2014)), which could also confound our analysis given that the strength of TE-mediated epigenetic effects vary along the same axis (**Figure 2**). Accordingly, we studied the associations between TE-induced H3K9me2 enrichment and TE population frequencies among *copies of the same TE family within species*, aiming to exclude the potential confounding effects of family identity on TE population frequencies. We performed logistic regression analysis for individual TE families that have at least ten insertions while accounting for the effects of TE length (population frequency ∼ epigenetic effects + length). With collinearity among predictor variables (see above) and the small number of TEs included for each TE family (fewer than 30 copies for 19 out 23 TE families, with a median of 19 of TEs included in the regression analysis), these analyses are again underpowered. For most of the TE families tested, we observed negative regression coefficient for the magnitude of TE-mediated H3K9me2 enrichment (**Table 2**), which is more than expected by chance (18 out of 23, *binomial test, p* = 0.0106). We observed similar results with the extent of TE-mediated H3K9me2 enrichment (17 out of 23, *binomial test, p* = 0.0346, **Table 2**). Despite the lack of statistical power, H and Cr1a families showed significant negative regression coefficient for TE-mediated epigenetic effects in one of the *D. yakuba* strains. Overall, our results revealed that, after controlling for the effects of TE length and family identity, we still found negative associations between the strength of TE-mediated epigenetic effects and TE population frequencies. By excluding the potential impacts of confounding factors on TE population frequencies, our observation strongly supports that TE-mediated enrichment of repressive marks is disfavored by natural selection in multiple species.

**Table 2.** Generalized linear regression coefficients for the effects of TE size and H3K9me2 enrichment (magnitude and extent) on the population frequencies of TEs from different families. Negative regression coefficients for TEs’ epigenetic effects on TE population frequencies are in bold. *p < 0.05.

| TE family | magnitude of H3K9me2 enrichment |  | extent of H3K9me2 enrichment |  |
| --- | --- | --- | --- | --- |
|  | TE size | magnitude | TE size | extent |
| ***D. melanogaster*** |  |  |  |  |
| BS | -8.10E-03 | **-2.65E+00** | -5.18E-03 | **-5.16E+02** |
| 297 | -4.86E-04 | **-1.43E+00** | -5.12E-04 | **-1.10E-04** |
| jockey | 2.17E-04 | **-1.30E+00** | 2.50E-04 | **-1.42E-03** |
| pogo | 9.46E-04 | **-1.04E+00** | 9.18E-04 | 9.47E-05 |
| Doc | 2.65E-04 | 2.20E-01 | 2.96E-04 | 2.55E-04 |
| hopper | -1.53E-04 | 1.36E+00 | -1.63E-04 | **-2.14E-04** |
| ***D. mauritiana*** |  |  |  |  |
| HB | -2.94E-03 | **-2.10E-01** | -2.96E-03 | **-3.23E-05** |
| Bari | -4.78E-03 | 6.58E-01 | -4.96E-03 | 3.65E-04 |
| hopper | 6.66E-03 | **-1.96E+00** | 8.70E-03 | **-3.72E-04** |
| ***D. simulans (strain 1)*** |  |  |  |  |
| H | 1.84E-02 | **-9.70E-02** | 1.76E-02 | **-3.08E-04** |
| transib2 | -3.69E-03 | **-1.26E+00** | -3.71E-03 | **-3.08E-04** |
| Tc1 | -1.19E-01 | **-3.56E-01** | -1.16E-01 | **-2.33E-04** |
| 1360 | -3.84E-02 | 1.37E-01 | -4.02E-02 | 8.88E-05 |
| diver2 | 2.16E-04 | **-1.39E-01** | -2.54E-04 | **-1.39E-03** |
| Helena | -3.99E-03 | 2.85E-01 | -3.62E-03 | 4.20E-04 |
| HB | -2.45E+00 | **-4.26E+00** | -2.48E+00 | **-1.61E-03** |
| roo | -4.01E-04 | **-2.71E-01** | -5.67E-04 | 4.10E-04 |
| ***D. simulans (strain 2)*** |  |  |  |  |
| H | -4.97E-03 | **-6.63E-01** | -4.94E-03 | **-3.02E-05** |
| Tc1 | -2.28E-03 | **-8.68E-01** | -2.46E-03 | **-7.59E-04** |
| ***D. yakuba (strain 1)*** |  |  |  |  |
| H | -1.80E-03 | **-3.79E-01** | -1.50E-03 | **-2.75E-04** |
| Cr1a | 9.53E-05 | **-6.66E-01** | -1.05E-04 | **-3.35E-04** |
| ***D. yakuba (strain 2)*** |  |  |  |  |
| H | -2.29E-02 | **-3.89E-01** | -5.48E-02 | **-8.98E-03\*** |
| Cr1a | 1.32E-03 | **-1.32E+00\*** | 4.42E-04 | **-1.27E-04** |

### Epigenetic effects of TEs negatively associate with genomic TE abundance across species

The magnitude and extent of TE-mediated H3K9me2 enrichment vary not only within genomes and between individuals, but also between species (**Figure 1**). According to a previously proposed hypothesis (Lee and Karpen 2017), this between-species difference in the strength of TE-mediated H3K9me2 enrichment could determine the average strength of selection removing TEs (**Figure 4**), leading to species-specific genomic TE abundance. To test this prediction, we investigated the associations between euchromatic TE numbers and the average magnitude and extent of TE-induced local enrichment of H3K9me2. Because the assignment of TEs into families is biased against TEs in the yakuba complex species (**Figure 2 – Supplementary Figure 3**), all TEs, irrespective of whether we could assign their family identity, were included in this between-species analysis (see Materials and Methods).

We found that TEs in species of the yakuba complex show a much larger magnitude of H3K9me2 enrichment than those in species of the melanogaster complex (**Figure 5A**, *Mann-Whitney U test, p* = 0.029). After controlling for the strong impacts of species complex, we found significant negative associations between the number of euchromatic TEs and the average magnitude of TE-mediated H3K9me2 enrichment across genomes (**Figure 5A**, TE abundance ∼ species complex + epigenetic effects; regression coefficient for the magnitude of H3K9me2 enrichment: -1603.5; ANOVA F-value: 9.05, df = 1, *p =* 0.030). On the other hand, the extent of TE-mediated H3K9me2 spreading does not differ between species complexes (*Mann-Whitney U test, p* = 0.89) nor associate with the number of euchromatic TEs (**Figure 5A**, regression coefficient for the extent of H3K9me2 enrichment (kb): 0.92; ANOVA F-value: 0.03, df = 1, *p* = 0.87, see Discussion). This finding echoes our observations of the nearly absent associations between the extent of TE-mediated H3K9me2 enrichment and TE population frequencies (**Figure 4**). To account for the phylogenetic non-independence among species, we used phylogenetic generalized least squares (PGLS, (Grafen 1989; Martins and Hansen 1997)) to repeat the regression analysis and found consistent results (regression coefficient for the magnitude of H3K9me2 enrichment: -1948; ANOVA F: 7.5, df = 1, *p =* 0.03; regression coefficient for the extent of H3K9me2 enrichment (kb): 599; ANOVA F: 0.14, df = 1, *p* = 0.72). Overall, the negative associations between the magnitude of TE-mediated H3K9me2 enrichment and abundance of euchromatic TEs support our prediction that varying strength TE-mediated epigenetic effects could drive between-species differences in genomic TE abundance.

**Figure 5.**
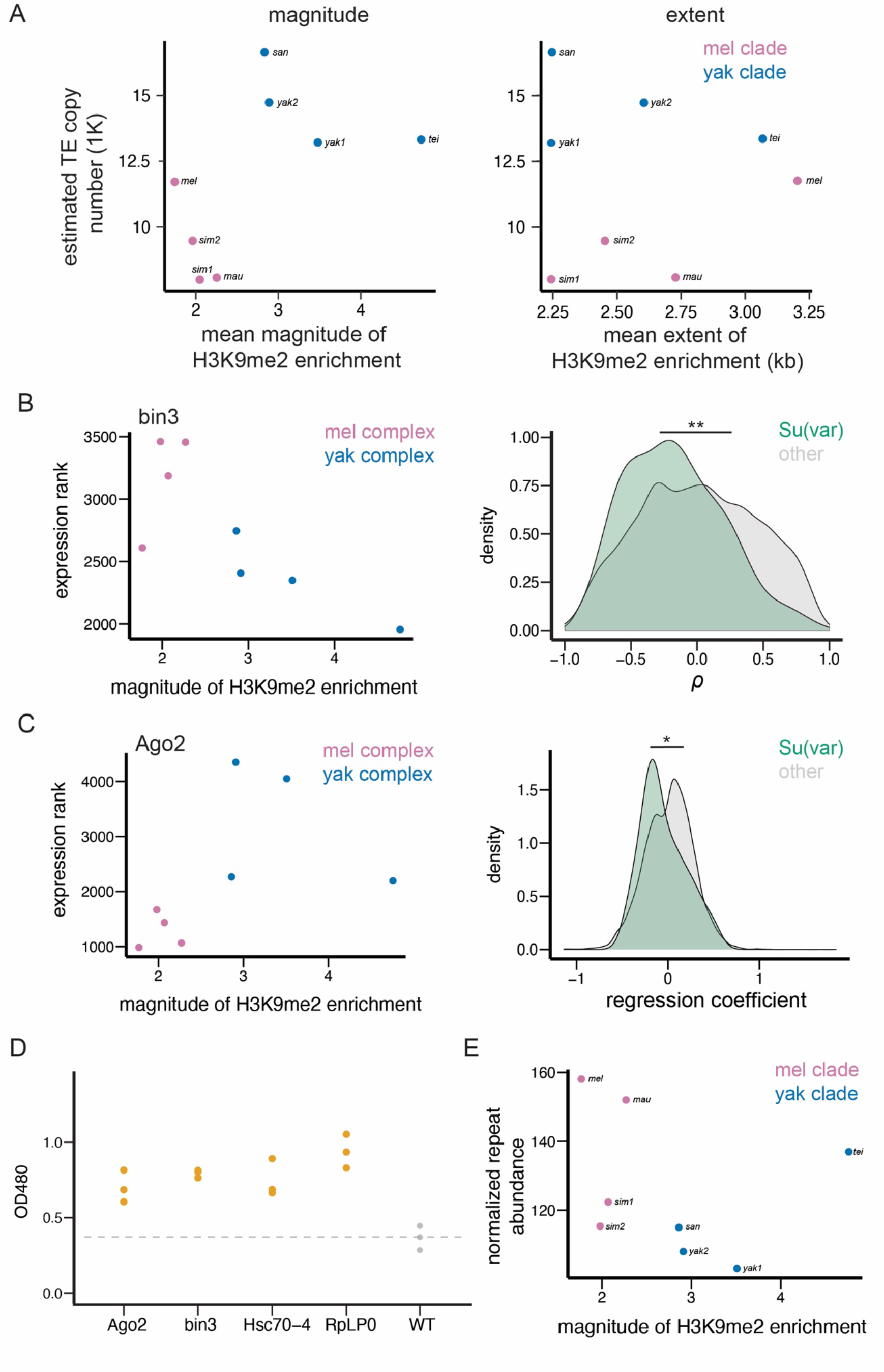
TE-mediated epigenetic effects associate with genomic TE abundance and repressive chromatin landscape. **(A)** The mean magnitude (left) and extent (right) of TE-mediate enrichment of H3K9me2 associate with the number of estimated euchromatic TEs across species. **(B)** An example gene (*bin3*) whose expression rank negatively correlates with the mean magnitude of TE-mediated H3K9me2 enrichment across species (left). The distributions of *Spearman rank correlation coefficient* (*⍴*) between the magnitude of TE-mediated H3K9me2 enrichment and the expression ranks significantly differ between *Su(var)* genes (green) and other genes in the genome (gray; right). **(C)** An example gene (*Ago2*) whose expression rank negatively correlates with the magnitude of TE-mediated H3K9me2 enrichment between species *within species complex,* but differs significantly between species complex (left). The distributions of *Spearman rank correlation coefficient* (*⍴*) for the effect of a gene’s expression on the magnitude of TE-mediated H3K9me2 enrichment significantly differs between *Su(var)* genes (green) and other genes in the genome (gray, right). (D) The reduced dosage of *Su(var)* genes influence the epigenetic silencing effect of *1360* TE. For each candidate *Su(var)*, we performed three replicates (three independent crosses) and one dot represents one cross. (E) The mean magnitude of TE-mediated enrichment of H3K9me2 associates with the abundance of H3K9me2-enriched Kmers. \*\**p <* 0.01 and *\*p <* 0.05 for *Mann-Whitney U test* or *Kolmogorov–Smirnov test* (see text).

### Species-specific repressive chromatin landscape associates with between-species differences in epigenetic effects of TEs

If varying strength of epigenetic effects of TEs indeed contributes to between-species differences in euchromatic TE copy number, as suggested by our observations (**Figure 5A**), species-specific differences in genetic factors that modulate the epigenetic effects of TEs could be the ultimate drivers for varying genomic TE abundance. Consistent with this possibility, we observed that the strength of TE-mediated epigenetic effects strongly depends on the host genetic background (**Figure 2**). Several candidate genetic factors were previously proposed to modulate the epigenetic effects of euchromatic TEs (Lee and Karpen 2017). These include *Su(var)* genes, whose protein products act as structural or enzymatic components of the heterochromatin (Girton and Johansen 2008; Elgin and Reuter 2013; Swenson *et al*. 2016), and heterochromatic repeats, which are targets of heterochromatin formation (Girton and Johansen 2008; Elgin and Reuter 2013). The ratio between the dosage of these two players (targets and regulators) was postulated to influence the nucleation and formation of constitutive heterochromatin and accordingly the spread of repressive marks from constitutive heterochromatin to euchromatic genome (Locke *et al*. 1988).

In order to test the hypothesized role of *Su(var)s* and heterochromatic repeats in shaping between-species differences in TE-mediated epigenetic effects, multi-species comparisons that span a reasonable phylogenetic resolution would be needed. Also, it would be important to demonstrate that *Su(var)s* that regulate the spreading of heterochromatic marks from constitutive heterochromatin play a similar role for the epigenetic effects of TEs in the euchromatic genome. This is because, though both enriched with H3K9me2/3, heterochromatin is comprised of large blocks of repetitive sequences enriched for repressive epigenetic marks (Riddle *et al*. 2011), whereas euchromatic TEs, though epigenetically silenced, are relatively short (Lee 2015; Lee and Karpen 2017) and usually surrounded by sequences enriched for active epigenetic marks (Kharchenko *et al*. 2011).

To test our hypothesis that species-specific differences in genetic factors contribute to varying epigenetic effects of TEs, we investigated whether there are positive associations between TE-mediated epigenetic effects and the dosage of *Su(var)* genes, which promotes the spreading of repressive marks from constitutive heterochromatin (reviewed in (Girton and Johansen 2008; Elgin and Reuter 2013)). We used the expression of *Su(var)* genes in 16-18hr embryos, a developmental stage matching our epigenomic data, as a proxy for the dosage of Su(var) protein products. Because we only found that the magnitude, but not the extent, of TE-mediated H3K9me2 enrichment negatively associates with TE copy number (**Figure 5A**), our following analysis focused on the magnitude of TEs’ epigenetic effects, with the aim of identifying the ultimate cause for varying genomic TE abundance. We estimated the *Spearman rank* correlation coefficients (*⍴*) between the expression rank of a gene (rank from the highest expressed genes) and the average magnitude of TE-mediated H3K9me2 enrichment across genomes (see **Figure 5B** for an example, also see Materials and Methods). A negative correlation indicates that a *Su(var)* gene has higher expression (and thus lower expression rank) in a genome with stronger epigenetic effects of TEs. We found that *Su(var)* genes, as a group, have significantly lower *⍴* than other genes in the genome (*Mann-Whitney U test, p =* 0.0073) and have a shifted distribution of *⍴* towards smaller value (*Kolmogorov–Smirnov test, p =* 0.033, **Figure 5B**). These observations suggest that the expression levels of *Su(var)s* correlate more positively with the magnitude of TE-mediated H3K9me2 enrichment than other genes in the genome. Among analyzed *Su(var)s*, the *⍴* for *bin3*, promoter of small-RNA mediated silencing (Singh *et al*. 2011), is among top 5% of all genes. *Lhr* and *HP4*, both of whose protein products are structural component of heterochromatin (Greil *et al*. 2007), and *Hsc70-4*, whose protein product is an interactor of core heterochromatin protein HP1a (Swenson *et al*. 2016), are among top 10% genome-wide (**Table S1**).

Intriguingly, the expression rank of *Ago2*, a key gene in initiating epigenetic silencing and a *Su(var)* (Deshpande *et al*. 2005), is among the top 10% genome-wide that *positively* correlates with TE-mediated enrichment of H3K9me2 (**Figure 5 – source data 1**), an association that is opposite to prediction. Upon further examination, we found that the expression of *Ago2* significantly differs between species in the two species complexes, which drives the positive correlation (**Figure 5C**). Accordingly, we performed regression analyses that associate the magnitude of TE-mediated H3K9me2 enrichment and gene expression rank while accounting for the effects of species complex (magnitude ∼ gene expression + complex). Similar to analysis based on *⍴* (seeabove, **Figure 5B**), the regression coefficients are significantly smaller for *Su(var)s* than those for other genes (*Mann-Whitney U test, p =* 0.063; *Kolmogorov–Smirnov test, p =* 0.032; **Figure 5C**), further corroborating our findings. With the regression analysis, we found that the association between *Ago2* expression rank and the magnitude of TE-mediated epigenetic effects is negative and among 10% genome-wide. We also found other *Su(var)* genes whose regression coefficients are among the top 10% of all genes, which includes *RpLP0*, whose protein product is core interactor of HP1a (Frolov and Birchler 1998), and *Su(var)3-3*, which codes for an eraser of active histone modification (Rudolph *et al*. 2007), **Figure 5 – source data 1**).

To confirm that these identified candidate *Su(var)s* also modulate TE-mediated local spreading of H3K9me2 in the euchromatic genomes, we leveraged a previously published reporter system that allows the quantification of TE-mediated epigenetic effects (Sentmanat and Elgin 2012). In this system, a DNA-based TE, *1360*, results in the enrichment of H3K9me2/3 at the adjacent *mini-white* reporter gene in the euchromatic genome, resulting in stochastic gene silencing and thus reduced amounts of red eye pigmentation (Sentmanat and Elgin 2012). By using existing loss-of-function *Su(var)* mutants, we found that reduced dosage (hemizygous) of candidate *Su(var)s* lead to significantly elevated levels of eye pigmentation, or reduced TE-mediated silencing effects (**Figure 5D**), supporting the roles of candidate *Su(var)s* in modulating the epigenetic effects of euchromatic TEs. With the assumption that the functional roles of *Su(var)* genes are conserved across *Drosophila* species studied, our findings suggest that the observed species-specific expression of *Su(var)s* could drive between-species differences in TE-mediated H3K9me2 enrichment in the euchromatic genome.

We also investigated whether the epigenetic effects of TEs negatively associate with the abundance of heterochromatic repeats, which was found to weaken the spreading of H3K9me2/3 from constitutive heterochromatin (reviewed in (Girton and Johansen 2008; Elgin and Reuter 2013)). We identified Kmers that are enriched with H3K9me2 in our ChIP-seq data and quantified their abundance using Illumina sequencing with PCR-free library preparation, in an effort to avoid biases in quantification of simple repeats (Wei *et al*. 2018) (see Material and Methods). Consistent with the prediction that repressive chromatin landscape weakens with increased abundance of heterochromatic repeats, we found that the magnitude of TE-mediated H3K9me2 enrichment is negatively associated with the abundance of H3K9me2-enriched repeats between species, though the comparison is not significant (**Figure 5E**, *Spearman rank correlation coefficient* (*⍴*) *= -0.595, p =* 0.13). Overall, our observations support the prediction that species-specific repressive chromatin landscape, which is shaped by the expression level of *Su(var)* genes and abundance of heterochromatic repeats, associates with differences in TE- mediated epigenetic effects between species.

## Discussion

The replicative nature of TEs have made them successful at occupying nearly all eukaryotic genomes. Yet, their “success” or genomic abundance drastically varies across the phylogenetic tree (Huang *et al*. 2012; Elliott and Gregory 2015; Wells and Feschotte 2020) and between closely related species (Hu *et al*. 2011; Rius *et al*. 2016; Legrand *et al*. 2019), raising important questions regarding the evolutionary causes and functional consequences of varying genomic TE abundance. Phylogenetic signals may explain some of the differences in TE abundance between distantly related taxa (Wells and Feschotte 2020) and, sometimes, between species within a taxa (Szitenberg *et al*. 2016). Variation in TE activities was also postulated to contribute to the wide variability of TE abundance ((Chen *et al*. 2019; Wong *et al*. 2019), but see (Ho *et al*. 2021)). Still another plausible cause is systematic differences in the strength of natural selection removing TEs between species. Investigations of this hypothesis have been largely focused on population genetic parameters that influence the efficacy of natural selection, such as mating systems (Wright and Schoen 1999; Dolgin and Charlesworth 2006; Boutin *et al*. 2012; Arunkumar *et al*. 2014; Ågren *et al*. 2014) and effective population size (Lynch and Conery 2003; Mérel *et al*. 2021). Here, our multi-species study tested step-by-step yet another evolutionary mechanism by which the strength of selection removing TEs could differ—through differences in the epigenetic effects of TEs that is driven by species-specific host chromatin landscape.

In this study, we investigated the prevalence, variability, and evolutionary importance of “the epigenetic effects of TEs”—TE-mediated local enrichment of repressive epigenetic marks—in the euchromatic genomes of six *Drosophila* species. These species come from two important and well-studied complexes within the *melanogaster* species subgroup (*melanogaster* and *yakuba* complexes), providing good phylogenetic resolution to decipher the evolutionary role of epigenetic mechanisms in shaping species-specific TE abundance. We observed that TE-mediated spreading of repressive marks is prevalent in all species studied and widely varies both within and between genomes. Interestingly, in addition to the intrinsic biological properties of TEs, our analyses revealed that host genetic background plays a critical role in determining the strength of TE-mediated epigenetic effects. These TE-mediated enrichments of repressive marks alter the epigenetic states of neighboring euchromatic sequences, including actively transcribing genes. Importantly, TEs exerting such epigenetic effects are selected against across multiple species, and the strength of this TE-mediated epigenetic effect negatively correlates with genomic TE abundance. Our findings extend previous studies based on few species and provide one of the first support for the importance of the inadvertent harmful effects of TE epigenetic silencing in shaping divergent genomic TE landscapes.

Intriguingly, while we found that these TE-mediated epigenetic effects increase the enrichment of repressive epigenetic marks at adjacent genes, we found limited evidence supporting that such effect reduces neighboring gene expression, echoing previous observations (Quadrana *et al*. 2016; Stuart *et al*. 2016; Lee and Karpen 2017). This lack of impacts of TE-mediated H3K9me2 enrichment on gene expression could have resulted from the complex relationship between repressive epigenetic modification and gene expression (de Wit *et al*. 2007; Yasuhara and Wakimoto 2008; Riddle *et al*. 2011; Meng *et al*. 2016; Caizzi *et al*. 2016), the varying sensitivity of genes to the enrichment of repressive marks (Rudolph *et al*. 2007; Vogel *et al*. 2009; Riddle *et al*. 2011), and the presence of other types of variants that also modulate gene expression (Stranger *et al*. 2007). Interestingly, our findings suggested another possibility—the transcription of genes, in return, influences TE-mediated epigenetic effects across the genome. Specifically, we found a more restricted spreading of repressive marks from TEs on the side facing a gene than the intergenic side, and this difference is only observed for TEs near highly expressed genes (**Figure 3**). It is worth noting that this difference in the extent of H3K9me2 spreading between the two sides of a TE is unlikely driven by the presence of insulator sequences (Gaszner and Felsenfeld 2006) because of the limited associations between the presence of insulator sequences and the reduction in TE-mediated H3K9me2 spreading on the gene side (**Figure 3 – Supplementary Figure S8**). On the other hand, active histone modifications enriched at transcriptionally active genes could antagonize the assembly of heterochromatin (Allshire and Madhani 2018), potentially restraining the spreading of repressive marks from silenced TEs. Consistently, in mice, active histone modification was reported to spread from an actively transcribing candidate gene into an adjacent TE (Rebollo *et al*. 2012). Similarly, active gene transcription at *D. miranda* neo-Y chromosome was found to impede the formation of heterochromatin (Wei *et al*. 2020). Future analysis on the distributions of active histone modifications may help reveal the mechanistic cause for our observed genome-wide dependencies of the extent of TE-mediated epigenetic effects on the transcriptional activities of adjacent euchromatic genes.

If not due to reducing the expression of adjacent genes, what functional consequences of TE-mediated epigenetic effects could have impaired host fitness and led to the observed selection against these TEs? Euchromatic TEs enriched with repressive epigenetic marks were reported to spatially interact with constitutive heterochromatin through phase separation mechanisms, a process observed to alter 3D structures of genomes and inferred to lower host fitness (Lee *et al*. 2020). In addition, the epigenetic effects of TEs could shift the usual DNA repair process in the gene-rich euchromatic genome, perturbing the maintenance of local genome integrity. This is because double stranded breaks happening in constitutive heterochromatin are repaired through a distinct cellular process from those in the euchromatic genome, owing to the enrichment of repressive epigenetic modifications (Chiolo *et al*. 2011; Janssen *et al*. 2016, 2019).

Still, the variance of gene expression was shown to be shaped by natural selection (Metzger *et al*. 2015; Duveau *et al*. 2018). Given the variegating properties of the spreading of repressive marks (reviewed in (Elgin and Reuter 2013)), TE-mediated epigenetic effects could have shifted the variance, instead of the mean, of neighboring gene expression, impacting host fitness.

Building upon our findings about the prevalence and variability of TE-mediated epigenetic effects and selection against such effects, our multi-species analysis provides strong support for the previously proposed role of TE-mediated epigenetic effects in determining genomic TE abundance (**Figure 6**). By combining transcriptomic analysis and *Drosophila* genetics experiment, we further revealed that between-species differences in TEs’ epigenetic effects could be driven by species-specific expression levels of host genetic factors that modulate heterochromatin. These findings connect the evolution of host genome (genomic TE abundance) with chromatin landscape through the inadvertent harmful effects of epigenetic silencing of TEs (**Figure 6**). It is worth noting that, within species complex, estimated nucleotide polymorphism, an indicator for effective population size (Charlesworth 2009), largely follows a similar rank order to our observed magnitude of TE-mediated H3K9me2 enrichment (nucleotide polymorphism– *melanogaster* complex: *D. simulans > D. mauritiana > D. melanogaster* (Langley *et al*. 2012; Meiklejohn *et al*. 2018) and *yakuba* complex: *D. tessieri > D. yakuba > D. santomea,* (Bachtrog *et al*. 2006); the only difference in the rank order being the epigenetic effect of TEs is stronger in *D. mauritiana* than in *D. simulans,* **Figure 5A**). Accordingly, varying efficacy of selection in purging deleterious TE insertions might also contribute to the observed between-species difference in TE abundance. Alternatively, sudden reduction in effective population size and accordingly weakened effectiveness of natural selection at removing TEs could drive the accumulation of TEs in some of the species (e.g., (Mérel *et al*. 2021)). Under this scenario, selection may favor epigenetic regulation that limits excessive inadvertent spreading of repressive marks from euchromatic TEs and could also result in the observed negative associations between TE-mediated epigenetic effects and genomic TE abundance.

**Figure 6.**
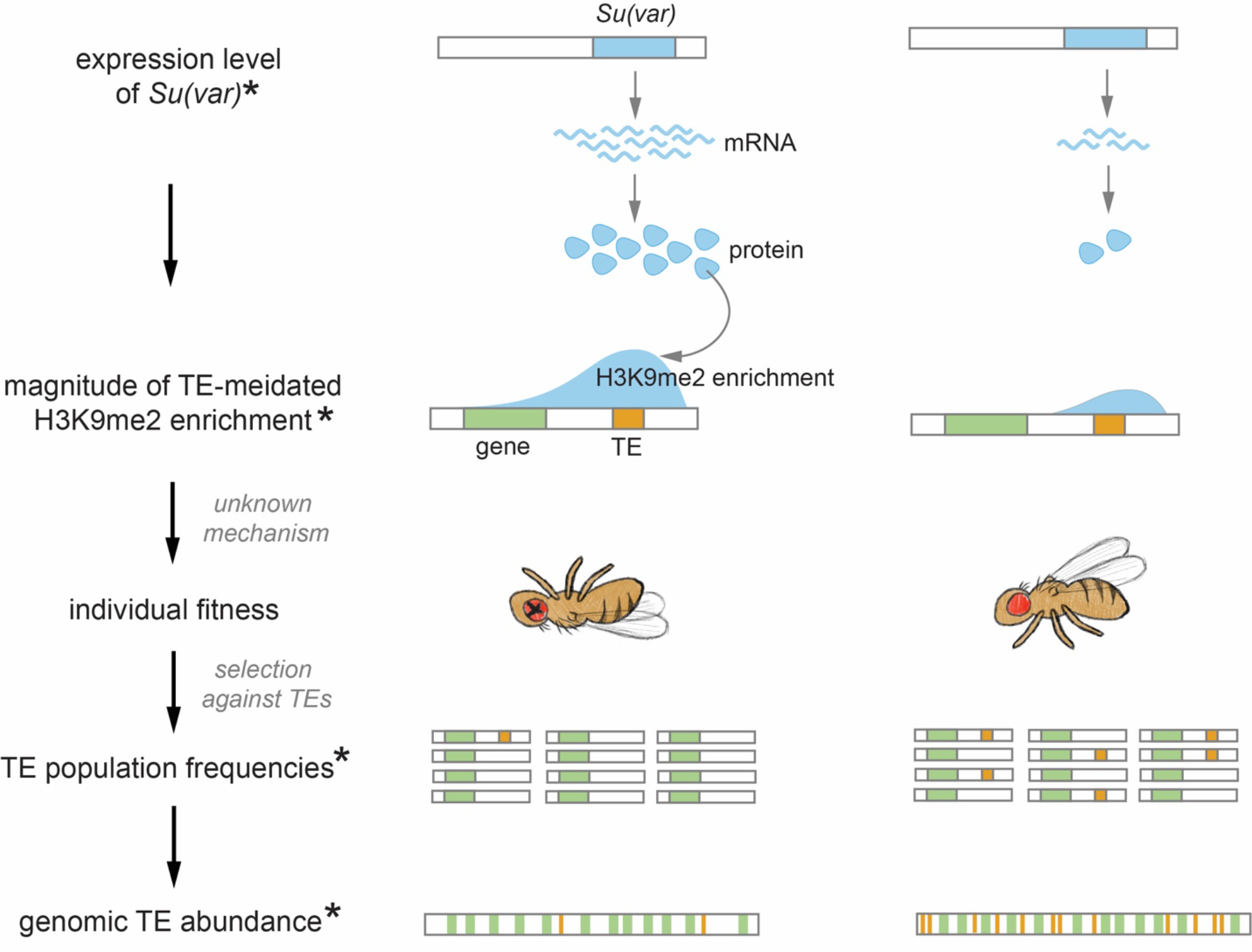
Proposed role of the chromatin landscape in determining genomic TE abundance. Our observations suggest that higher expression of *Su(var)s,* which promote repressive chromatin environment, would result in stronger TE-mediated epigenetic effects (e.g., species on the left). With mechanisms that are yet to be revealed (see Discussion), the stronger epigenetic effects of TEs will reduce individual fitness and, accordingly, stronger selection against TEs, driving lower population frequencies of TEs and an overall lower genomic TE abundance. Observations made in this study are denoted with *.

Curiously, we only found the predicted negative association between genomic TE abundance and the *magnitude* of TE-mediated H3K9me2 enrichment across species, but not the *extent* of the spreading, even though these two indexes of a TE significantly correlate within genomes (**Figure1 – supplementary Figure S1**). Similarly, the evidence for selection against TE-mediated epigenetic effects is stronger for the magnitude of TE-induced H3K9me2 enrichment than for the extent (**Figure 4**). Previous empirical (Talbert and Henikoff 2000) and theoretical (Erdel and Greene 2016) studies have suggested that the spreading of repressive heterochromatic marks is a nonlinear process. Accordingly, the extent of the spreading of repressive marks from TEs could be sensitive to the genomic context and subject to substantial stochasticity. This echoes our observations that the transcription of genes influences the extent of TE-mediated H3K9me2 spreading, but not the magnitude of H3K9me2 enrichment (**Figure 3D and Figure 3 – Supplementary Figure S5**). The magnitude of TE-mediated H3K9me2 enrichment, which is estimated in windows right next to TE boundary, may thus be a more direct measurement for the strength of TE-mediated epigenetic effects and more indicative of the associated harmful impacts.

Proper maintenance of heterochromatin ensures genome function and integrity (reviewed in (Janssen *et al*. 2018)). Intriguingly, the expression level (see above), copy number (Levine *et al*. 2012; Ross *et al*. 2013; Helleu and Levine 2018), and amino acid sequences (Levine *et al*. 2012; Sasaki *et al*. 2019) of heterochromatin structural and enzymatic components have been observed to vary substantially within and between species. The abundance and composition of heterochromatic repeats also rapidly turnover between genomes (Larracuente 2014; Wei *et al*. 2014, 2018). While variation in these genetic factors were postulated to contribute to the evolution of heterochromatin functions (e.g., (Ferree and Barbash 2009; Helleu and Levine 2018; Mills *et al*. 2019)), our discoveries further extend their evolutionary impacts from the gene-poor genomic dark matter into the gene-rich euchromatin. Heterochromatin modulators may not only determine TEs’ local epigenetic impacts on flanking sequences, but also shape TE abundance and, accordingly, the structure and function of the euchromatic genomes. The evolution of euchromatin and heterochromatin, two genomic compartments usually presumed to function independently, may be more interconnected than previously thought by the inadvertent deleterious epigenetic effects of a widespread genetic parasite.

## Materials and Methods

### Drosophila strains

Drosophila strains used in this study include *D. melanogaster* ORw1118 (Sexton *et al*. 2012), *D. simulans w501* (Drosophila Species Stock Center, strain 1) and *Mod6* (strain 2)*, D. mauritiana w12, D. yakuba* NY73PB (strain 1) and Tai 18E2 (strain 2), *D. santomea* AG01482, and *D. teissieri GT53W gen14*. Flies were cultured on standard medium at 25°C and 12hr/12hr light/dark cycles.

### ChIP-seq and RNA-seq experiment

We collected 16-18hr embryos from each strain to perform ChIP-seq and RNA-seq experiments (with two biological replicates). Before embryo collection, young adults (3-7 days old) were allowed to lay eggs on fresh apple juice plates for one hour at 25°C. We then collected embryos for two hours on another fresh apple juice plates and stored those collection plates at 25°C to enrich for 16-18hr embryos. Chromatin isolation and immunoprecipitation were performed following the modEncode protocol (http://www.modencode.org/) with the following modification. Before splitting the Input and IP samples, we added 4 ul of SNAP-ChIP K-MetStat Panel (EpiCypher) to every 10 ug of chromatin. This concentration allowed ∼200-500 barcoded H3K9me2 nucleosome reads with ∼15 million Illumina reads, which is the targeted sequencing depth of our samples. The SNAP-ChIP K-MetStat Panel serves as a spike-in and allows normalization between samples. We used H3K9me2 antibody (abcam 1220), which was validated by the modEncode and shows H3K9me2-specific binding (Egelhofer *et al*. 2011), to perform the ChIP experiment. ChIP-seq libraries were prepared using NEBNext Ultra DNA Library Prep Kit for Illumina (NEB) following manufacture’s protocol. RNAs were extracted using the RNeasy Plus kit (Qiagen) and library prep with Illumina TrueSeq mRNA stranded kit (Illumina) following manufacture’s protocols. Both ChIP-seq and RNA-seq samples were sequenced on Illumina Hi-Seq4000 with 100bp, paired-end reads.

### PacBio assemblies

We generated PacBio assemblies for *D. melanogaster* ORw1118 strain with following procedures. High molecular weight DNA was extracted from 450 adults (mixed sexes) using Qiagen Genomic-tip kit following manufacture’s protocol. Extracted DNA then underwent SMRTbell library preparation with target insert size as 20kb and sequenced on one SMRTcell using P6-C4 chemistry on a Pacific Biosciences RSII platform. We used Canu (version 2.0. (Koren *et al*. 2017)) to assemble a draft genome from the PacBio reads. We then used pbmm2 SMRT Analysis (version 7.0.0) to align the PacBio raw reads to the genome and used Arrow, also from SMRT Analysis, to polish the genome. The final genome has N50 = 12.1MB and 178MB for the total genome size.

PacBio assemblies for *D. simulans* and *D. mauritiana* are from (Chakraborty *et al*. 2021) and for D. yakuba (NCBI: GCA_016746365.2 and GCA_016746335.2), D. santomea (NCBI: GCA_016746245.2) and D. teissieri (GCA_016746235.2) are from the Andolfatto lab (at Columbia University ).

### Annotation of TEs in PacBio assemblies

We ran Repeatmodeler (version 2.0.1; (Flynn *et al*. 2020)) on PacBio genome assemblies. NCBI BLASTDB and the analyzed genomes were used as inputs for the BuildDatabase function in Repeatmodeler. We ran Repeatmodeler with options “-engine ncbi -LTRStruct” and selected the output repeats with class as LTR, LINE, DNA or unknown as potential TE sequences. We required a TE to be at least 500bp to be included in our analyses.

To assign family identity to identified TEs, we used an iterated blast approach. We first blasted the TE sequences (using blastn (Camacho *et al*. 2009)) to TEs annotated in *D. melanogaster* reference genome (version 6.32) and ‘canonical’ TE sequences of *Drosophila* (retrieved from Flybase Sep 2019) with following parameters: -evalue 1e-5; - perc_identity 80 for the melanogaster complex and -perc_identity 60 for the yakuba complex. When an identified TE has blast hits to multiple TE families, we only assigned the TE to a family when at least 80% of the covered query belongs to one and only one family. We then added the annotated TEs to the “blast database” and repeat the process three more times in order to allow the identification of TEs that are diverged from those in the reference *D. melanogaster* genome or canonical TEs annotated in various *Drosophila* species. TE insertions that are within 500bp were merged if they are from the same TE family or excluded from the analysis if they belonged to different families. We excluded DINE-1, which are mostly fixed in the *melanogaster* complex species (Kapitonov and Jurka 2003) but underwent a recent burst of activities in *D. yakuba* and likely other closely related species (Yang and Barbash 2008). We also excluded telomeric TEs (HeT-A, TART, and TAHRE), which predominantly locate at the end of chromosomes that are largely heterochromatic. Because most of the TEs included in our blast database are from *D. melanogaster,* it is plausible that our family assignment process is biased against assigning family identity to TEs in the species of the yakuba complex. Accordingly, except for analyses that require TE family identity (e.g., **Figure 2B and Figure 2C** and the estimation of TE population frequencies, see below), we included all TEs that are at least 500bp in the analysis, irrespective whether we could assign their family identities or not.

### Identification of Euchromatin/heterochromatin boundaries

Because we are interested in the epigenetic effects of euchromatic TEs, our analysis excluded those that are in or close to heterochromatin. To determine the euchromatin-heterochromatin boundaries in each genome, we generated H3K9me2 fold-enrichment tracks using Macs2 (Li et al. 2011). We then visualized the H3K9me2 enrichment genome-wide using IGV (version 2.10.2, (Thorvaldsdóttir *et al*. 2013)) to identify the sharp transition in H3K9me2 enrichment and used positions that are 0.5Mb “inward” (towards the euchromatin) from the transition as the euchromatin-heterochromatin boundaries. These conservative euchromatin-heterochromatin boundaries are expected to minimize the influence of consecutive heterochromatin in influencing our analysis.

### ChIP-Seq analysis

In order to align reads originated from the spike-in control (SNAP-ChIP K-MetStat Panel, Epicypher), we combined PacBio genomes with DNA sequence barcode of the SNAP-ChIP K-MetStat Panel, which were serve as the “reference genome” for Illumina sequence alignment. Raw reads were trimmed using *Trimmomatic* v 0.35 (Bolger *et al*. 2014) before aligning to the reference genome with bwa mem v 0.7.16a (Li and Durbin 2009). Read with low mapping quality (q < 30) and reads that mapped to multiple locations were removed using Samtools (Li 2011). We used the abundance of reads corresponding to the spike-in H3K9me2 nucleosome to normalize the background level of H3K9me2, following (Lam *et al*. 2019). Briefly, we calculated the enrichment of spike-in H3K9me2 nucleosome reads as: *Esi* = (barcode fragments in ChIP) / (barcode fragments in input). We then used bedtools v.2.25.0 (Quinlan and Hall 2010) to obtain read coverage across the genome for ChIP and Input samples. For 25bp nonoverlapping windows across the genome, we calculated the per-locus enrichment as *Elocus* = (fragment coverage in ChIP) / (fragment coverage in input). The histone modification density (HMD) for H3K9me2 in a particular 25bp window was then estimated as *Elocus*/*Esi*.

### Estimation of TE-mediated H3K9me2 enrichment

For each TE, we normalized the local H3K9me2 HMD level in its flanking sequence by the median HMD for regions 20-40kb upstream and downstream of each TE, following (Lee and Karpen 2017). The reasons behind this approach come from the observations that TE-mediated spreading of repressive epigenetic marks is usually within 10kb in *Drosophila* (Lee 2015; Lee and Karpen 2017). We then divided the 20kb upstream and downstream from a TE into 1kb nonoverlapping windows and, for each window, calculated the median of normalized H3K9me2 HMD among its 40 25bp-HMD units (see above, m-HMD). To estimate the magnitude of TE-mediated local enrichment of H3K9me2, we calculated the m-HMD for the 1kb left and right flanking regions separately and took the average of the two sides. To estimate the extent of H3K9me2 spreading, we examined whether m-HMD is above one, which indicates that the H3K9me2 HMD level for the window is higher than that of the local background. We scanned across windows, starting from those right next to TEs, and identified the farthest window in which the m-HMD was consecutively above one. We then used the average for the two sides as the extent of H3K9me2 spreading. The estimates for HMD magnitude or extent from two replicates positively correlate (*Spearman rank correlation coefficients* (*⍴*) = 0.19-0.84, *p <* 10^-8^, **Figure 1 – Supplementary Figure S4**). We thus averaged the estimates from two replicates to generate the magnitude and extent of H3K9me2 enrichment used in the analysis. It is worth noting that the estimated extent of H3K9me2 spreading with different thresholds of m-HMD strongly correlate (*Spearman rank correlation coefficients* (*⍴*) = 0.69-0.87, *p <* 10^-16^, **Figure 1 – Supplementary Figure S5)**, suggesting the robustness of such estimate. In the analysis, a TE was considered showing “epigenetic effects” when the magnitude of H3K9me2 enrichment is above one or has at least 1kb spreading of H3K9me2 (see text).

For two species (*D. simulans* and *D. yakuba*), we also investigated the effects of TEs by comparing the enrichment of H3K9me2 at homologous sequences with and without TE insertions and compare that to single species-based method (see above). We first ran MACS v 2.7.15 (Zhang *et al*. 2008) to identify shared peaks of H3K9me2 enrichment using liberal significant threshold (with broad-cutoff *p* = 0.5). TEs in these shared peaks were then excluded from the analysis because we could not determine if the enrichment of H3K9me2 for these regions were induced by TEs. We also excluded focal TEs whose homologous sequences were within 1kb of another TE in the alternative strain. The magnitude of H3K9me2 enrichment was estimated as the m-HMD of the focal TE in the focal strain standardized by the m-HMD in the 1kb TE-flanking regions in the homologous sequence in the alternative strain. The extent of H3K9me2 spreading was measured as the distance for the farthest windows from TEs that the m-HMD was consecutively higher in focal strain than that in the alternative strain. The estimates based on one or two genomes significantly correlate (**Figure 1 – Supplementary Figure S2**; see text).

### Inference of genic H3K9me2 enrichment

Genic H3K9me2 enrichment is estimated as the average of HMD of the gene body, excluding genes with TEs inserted. When comparing the H3K9me2 enrichment level of homologous genic alleles with and without adjacent TEs, we calculated z-score as: (mean HMD of allele with nearby TE - mean HMD of allele without nearby TE in the alternative strain)/(standard deviation of both strains).

### Association between TE-mediated epigenetic effects and TE abundance across species

To examine whether the epigenetic effects of TEs associate with TE abundance across species while accounting for species complex effects, we performed linear regression analyses using the following model: TE abundance ∼ species complex (*melanogaster* or *yakuba* complex) + epigenetic effect. ANOVA F-test was used to examine whether the epigenetic effect was significant. We also performed phylogenetic generalized least squares (PGLS, (Grafen 1989; Martins and Hansen 1997)) analysis using a tree adapted from (Turissini and Matute 2017; Chakraborty *et al*. 2019) with arbitrary branch lengths: ((((Dsim_strain1:0.1,Dsim_strain2:0.1):0.15, Dmau:0.25):3,Dmel:3.25):7.25, (((Dyak_strain1:0.1,Dyak_strain2:0.1):0.9,Dsan:1):1.75, Dtei:2.75):7.75). Although the significance level was sensitive to the within-species branch lengths, the sign of the coefficients remained unchanged. To fit the above model, we used gls function from nlme package with a Brownian correlation structure based on the tree (imported using ape package) in R.

### Gene expression analysis

Reference genome sequences and annotations for *D. melanogaster, D. simulans, D. mauritiana* and *D. yakuba* were downloaded from NCBI Datasets (*D. melanogaster*, version Release 6 plus ISO1 MT; *D.simulans*, version ASM75419v2; *D. mauritiana*, version ASM438214v1; *D. yakuba*, version dyak_caf1). The annotations were lifted to PacBio assemblies by aligning the PacBio assemblies to NCBI references using minimap v2.17 (Li 2018). Because the annotations for *D. santomea* and *D. teissieri* were not available when we performed the analyses, we used MAKER v2.31.8 (Holt and Yandell 2011 p. 2) to annotate the PacBio assemblies. Specifically, we used Trinity- v2.85 (Grabherr *et al*. 2011) to de novo assemble the mRNA-seq for the genome as EST evidence for MAKER. We also supplied the protein sequences from *D. melanogaster, D. sechellia, D. simulans, D. mauritiana, D. erecta* and *D. yakuba* as protein homology evidence for MAKER. We then ran MAKER with the default settings to obtain annotations for *D. santomea* and *D. teissieri.* To estimate gene expression abundance, we mapped the raw RNAseq reads to the annotated genomes with STAR v2.6.0 (Dobin *et al*. 2013) with options --quantMode TranscriptomeSAM GeneCounts -- chimFilter None. In order to compare the expression levels between different species, we used reciprocal best blasts between *D. melanogaster* and one other species to identify one-to-one orthologs. We then obtained a set of shared orthologs among six species to compare expression levels. For this set of shared orthologs, we estimated the RPKM (reads per kilobase per million reads) as the averaged RPKMs from two replicates (RPKMs from two replicates strongly correlate; *Spearman rank correlation coefficients* (*⍴*) > 0.98 for all genomes, *p <* 10^-22^). We then genes from the highest to lowest RPKMs in each strain to get expression rank.

### Estimation of TE population frequencies

Raw Illumina reads for *D. melanogaster* (Lack *et al*. 2015)*, D. simulans* (Rogers *et al*. 2014)*, D. mauritiana* (Garrigan *et al*. 2012), and *D. yakuba* (Rogers *et al*. 2014) were downloaded from SRA (SRP006733, SRP040290, SRP012053 and SRP029453 respectively). We used FastQC v0.11.7 (https://qubeshub.org/resources/fastqc) to check the read quality and TrimGalore v0.6.0 (“Babraham Bioinformatics - Trim Galore!”) to remove adapter and low-quality sequences. Illumina reads were then mapped to the corresponding PacBio assembly using bwa mem v 0.7.16a (pair-end mode). We further used samtools to filter out reads with mapping quality smaller than 50 (MAPQ < 50).

To call the presence/absence of TEs, we followed the basic ideas developed in (Cridland *et al*. 2013; Lee and Karpen 2017) with following modifications. Briefly, we parsed out reads that uniquely mapped to the +/-500bp around TEs annotated in the PacBio assembly using *seqtk v1.3-r107-dirty* (https://github.com/lh3/seqtk). Parsed reads were assembled into contigs using *phrap v1.090518* (Ewing and Green 1998) with parameters from (Cridland *et al*. 2013). We mapped the resultant contig to PacBio assembly using bwa mem (default options). If a *single* contig mapped across 20bp upstream and downstream of an annotated TE, a TE is called absent. To determine whether a TE is present, we evaluated whether *two* separate contigs spanned across the start and end of a TE’s boundaries respectively, with at least 30bp inside the TE and 20bp outside the TE and a minimum total alignment length of 50bp. We also evaluated contigs that mapped within annotated TEs. A TE is called present if there are two contigs spanning both the start/end of the TE insertion respectively *or* there is one contig spanning either the start or end of the TE and one contig aligned within TE insertion. A TE is considered missing data if none of the above criteria were met.

In the initial runs, we noticed that the failure to identify some TEs in the tested strain is due to the imprecise TE boundaries annotated by RepeatModeler 2. In order to reduce the rates of missing data, we refined the called TE boundaries with following procedures. TE absence calls could be viewed as deletion structural variants (SVs) with respect to the PacBio genome. We thus used *Lumpy-sv v0.2.13* (Layer *et al*. 2014) to call SVs in each strain. Deletions that overlap with the annotated TEs were extracted using sytyper v0.0.4 (Chiang *et al*. 2015) and merged using svtools v 0.5.1 (Larson *et al*. 2019) to refine TE boundaries, which were then used in the above pipeline. For most TEs, the differences between the updated and RepeatModeler 2 boundaries are short and only 5.2-7.9% of TEs boundaries in each genome were significantly updated (>= 20bp from original boundaries). Code for the TE calling pipeline can be found at https://github.com/harsh-shukla/TE_freq_analysis.

### Quantification of the heterochromatic repeats

To identify the repeat sequences enriched in the heterochromatic regions of the genome, we first used KMC3 (version 3.1.1, (Kokot *et al*. 2017 p. 3)) to quantify the 12- mers in our H3K9me2 IP and matching Input samples. In order to compare 12-mers abundance between IP and Input libraries, we normalized the 12-mers counts by the number of reads mapped uniquely to the PacBio genome with at least 30 mapping quality score. 12-mers that have at least a 3-fold enrichment in an IP sample when compared to its matching Input sample were considered as heterochromatic repeats.

Because PCR amplification of sequencing libraries were shown to influence the quantification of simple repeats (Wei *et al*. 2018), we sequenced the genomes of focused strains with Illumina PCR-free library preparation. We extracted DNA with 40 females using Qiagen DNeasy Blood & Tissue Kit, following manufacture protocol. Extracted DNA was then prepared into Illumina sequencing libraries with PCR-free protocol and sequenced with 150pb paired-end reads by Novogene (Sacramento, CA). We then ran KMC3 on these libraries to quantify the abundance of heterochromatic repeats identified above. In order to compare across strains, the number of heterochromatic repeats were further normalized with the total number of reads from either the orthologous region.

### Drosophila mutant crosses and eye pigmentation assay

To investigate the mutant effects of identified candidate *Su(var)s*, we followed approaches outlined in (Sentmanat and Elgin 2012). Specifically, we used a strain developed in (Sentmanat and Elgin 2012), which has a TE *1360* placed next to *mini-white in y; w* background. It was found that 1*360*-mediated epigenetic effects influence the expression level of *mini-white* and thus the intensity of eye pigmentation. Our analysis only tested the effects of *Su(var)s* whose existing mutants do not have any eye markers to avoid confounding effects on the quantification of eye pigmentation. We crossed 3-5 days old virgin females of this strain to males of the *Su(var)* mutant strains or a control strain and then quantified the eye pigmentation level in 25-30 3-5 days old F1 males following methods described in (Sentmanat and Elgin 2012). Three independent crosses were performed for each mutant. Strains used in the analysis include BDSC 6398 (*Hsc70-4* mutant), BDSC 11537 (*RpLP0* mutant), BDSC 30640 (*Bin 3* mutant) BDSC 36511 (*Ago 2* mutant), and BDSC 6559 (*y; w*, control).

## Acknowledgements

We greatly appreciate J.J. Emerson and Peter Andolfatto for generously providing the PacBio genomes and corresponding *Drosophila* strains, Sally Elgin for providing 1360- mini-white strains, Giacomo Cavalli for sharing ORw1118 line, Bloomington Drosophila Stock Center for providing *Su(var)* mutants, and David Acevedo and Jasmine Osei-Enin for technical assistance. Patrick Reilly and Mahul Chakraborty provided helpful guidance on the PacBio genome assembly, and Kevin Brick (from Camerini-Otero group) for discussion about ChIP-seq normalization. We thank University of California High-Throughput Genomics Facility and High Performance Cluster at UC Irvine for sequencing and computational resources. We also appreciate Aneil Agrawal, Ching-Ho Chang, Jae Choi, Brandon Gaut, Mia Levine, Aline Muyle and members of the Lee lab for providing helpful discussions and comments on the manuscript and Leila Lin for assistance in drawing the model figure. This work was supported by NIH R00GM121868 and R35GM14292 to YCGL.

## Notes

### Competing Interest Statement

The authors have declared no competing interest.

