## Supplemental Figures and Table for "Species-specific chromatin landscape determines how transposable elements shape genome evolution"

**Figure 1 – Supplementary Figure S1.** The magnitude and extent of TE-mediated H3K9me2 enrichment strongly correlate in all genomes studied. *Spearman correlation coefficients ($\rho$)* are all significantly different from 0 ($p$ < 2.2e$^{-16}$).

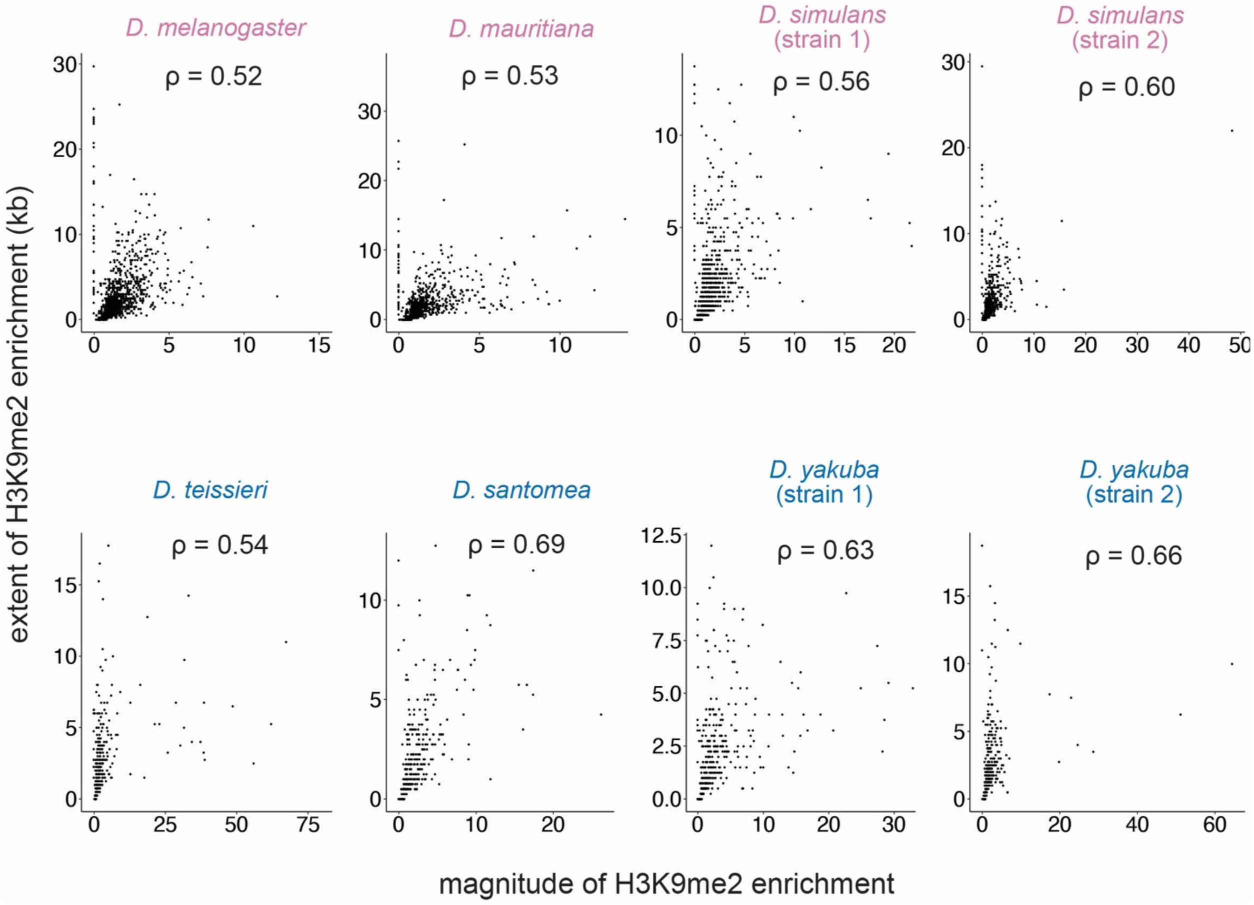

**Figure S1 – Supplementary Figure S2.** Estimates for the magnitude and extent of TE-mediated H3K9me2 enrichment that are based on one genome (x-axis) or two genomes (y-axis) strongly correlate. *Spearman correlation coefficients (ρ)* are all significantly different from 0 ($p < 10^{-10}$).

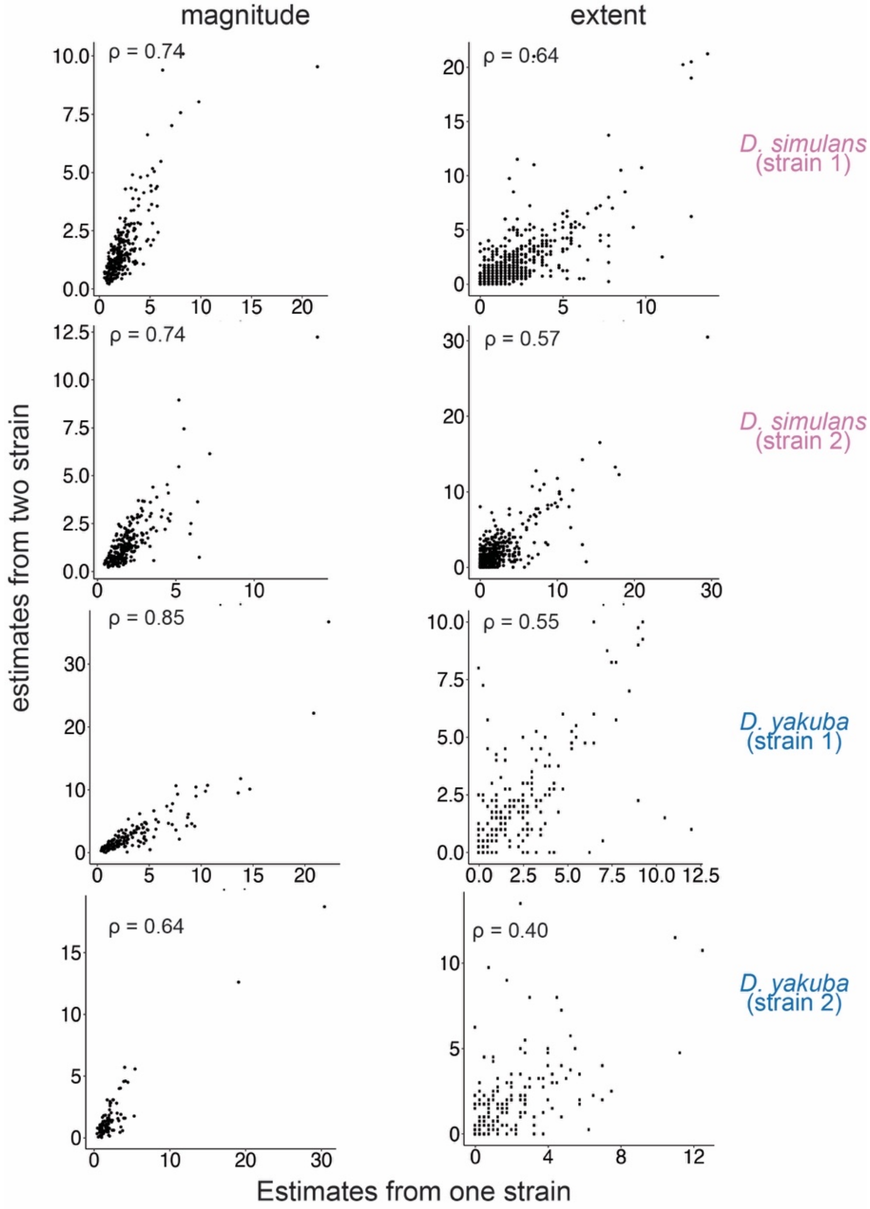

**Figure 1 – Supploementary Figure S3**. The magnitude/extent of TE-mediated H3K9me2 enrichment and TE length weakly correlate in most species studied. The *Spearman correlation coefficients (ρ)* are significantly different from zero at ***p* < 0.01, and **p* < 0.05.

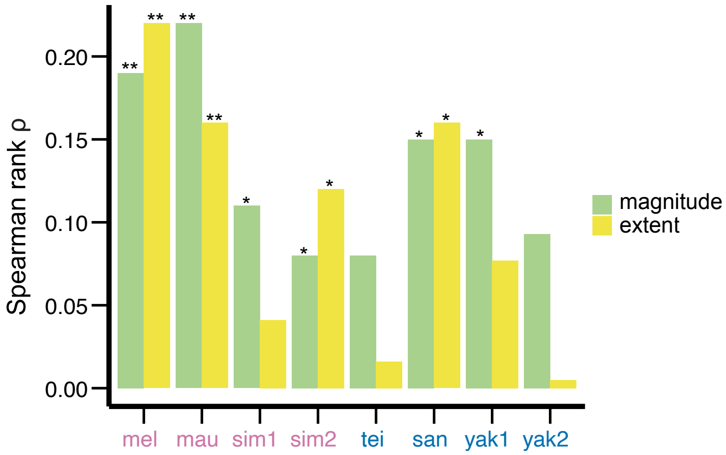

**Figure 1 – Supplementary Figure S4**. Estimates for the magnitude and extent of TE-mediated H3K9me2 enrichment significantly correlate between replicates. *Spearman rank correlation coefficients* ($\rho$) are all significantly different from 0 ($p < 10^{-8}$).

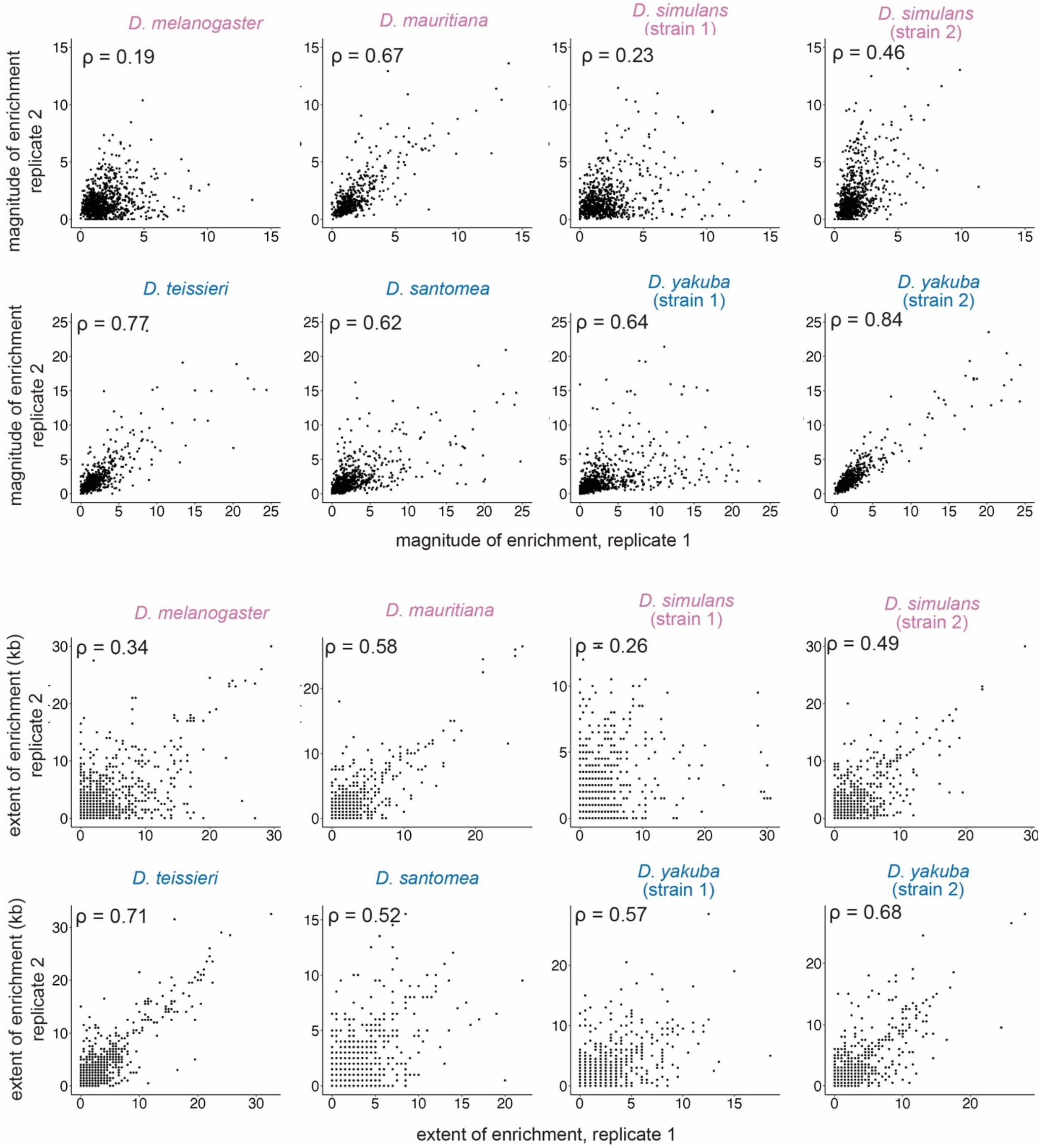

**Figure 1 – Supplementary Figure S5**. Estimates for the extent of TE-mediated H3K9me2 enrichment strongly correlate with different HMD cutoffs—HMD > 1 (used threshold through out the study), HMD > 1.5, and HMD > 2. *Spearman rank correlation coefficients* ($\rho$) are all significantly different from 0 ($p < 10^{-16}$).

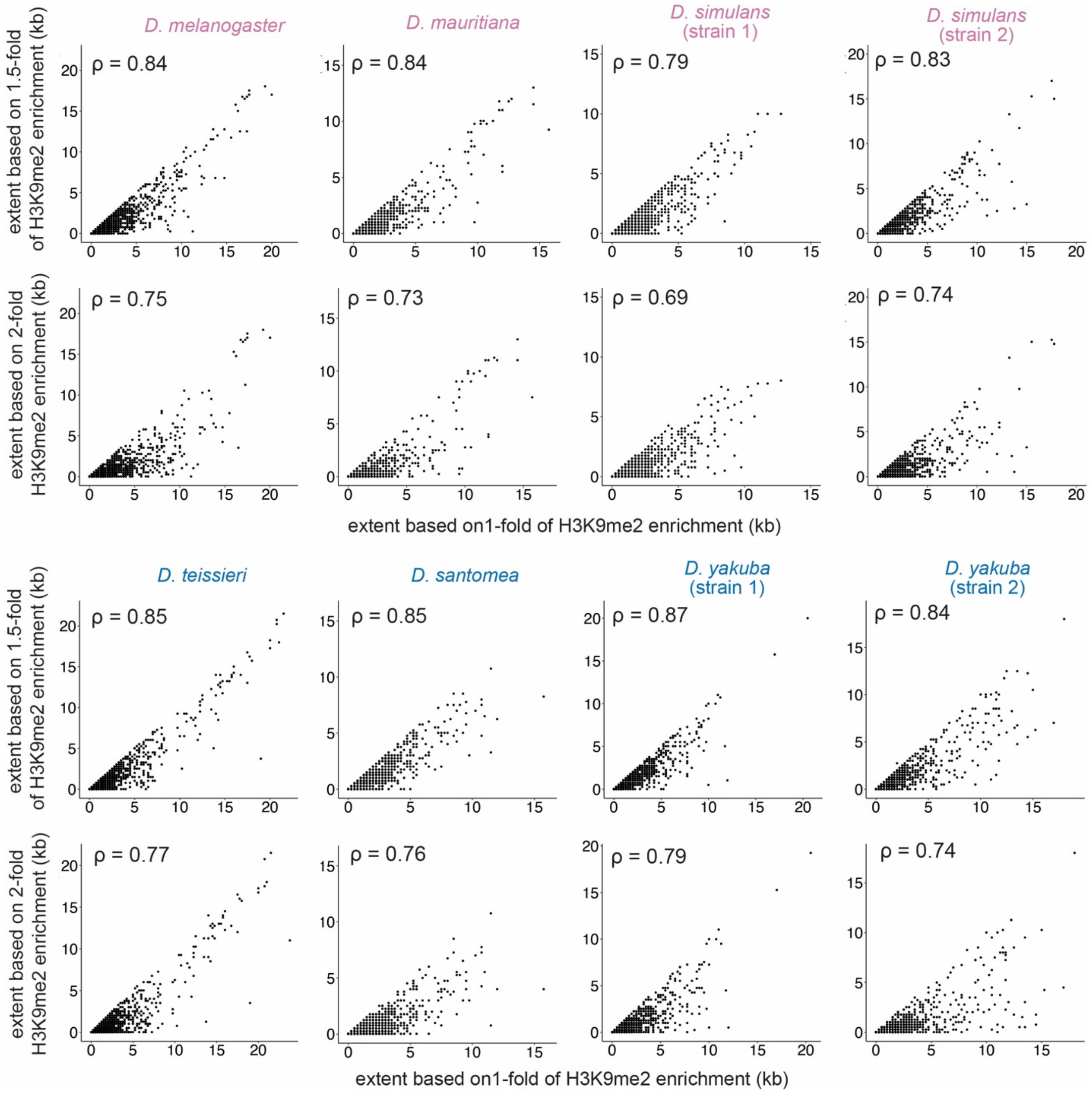

**Figure 2 – Supplementary Figure S1.** Magnitude and extent of TE-mediated H3K9me2 enrichment of different families for other *melanogaster* complex species (*D. melanogaster* and *D. mauritiana*).

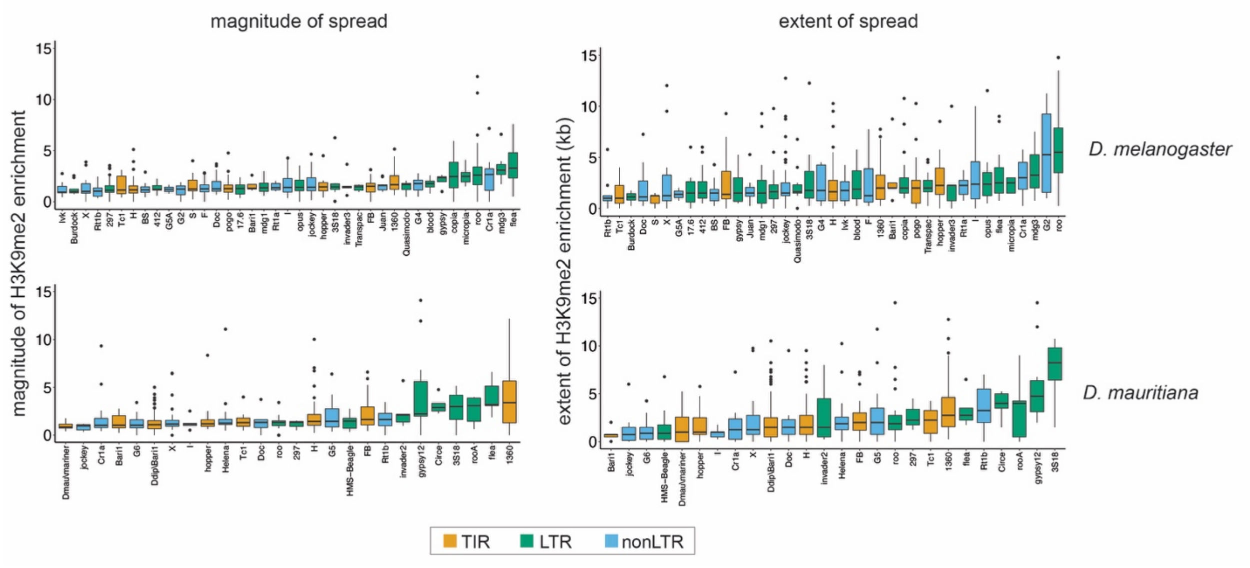

**Figure 2 – Supplementary Figure S2.** Magnitude and extent of TE-mediated H3K9me2 enrichment of different families for *yakuba* complex species (*D. teissieri, D. santomea,* and *D. yakuba*).

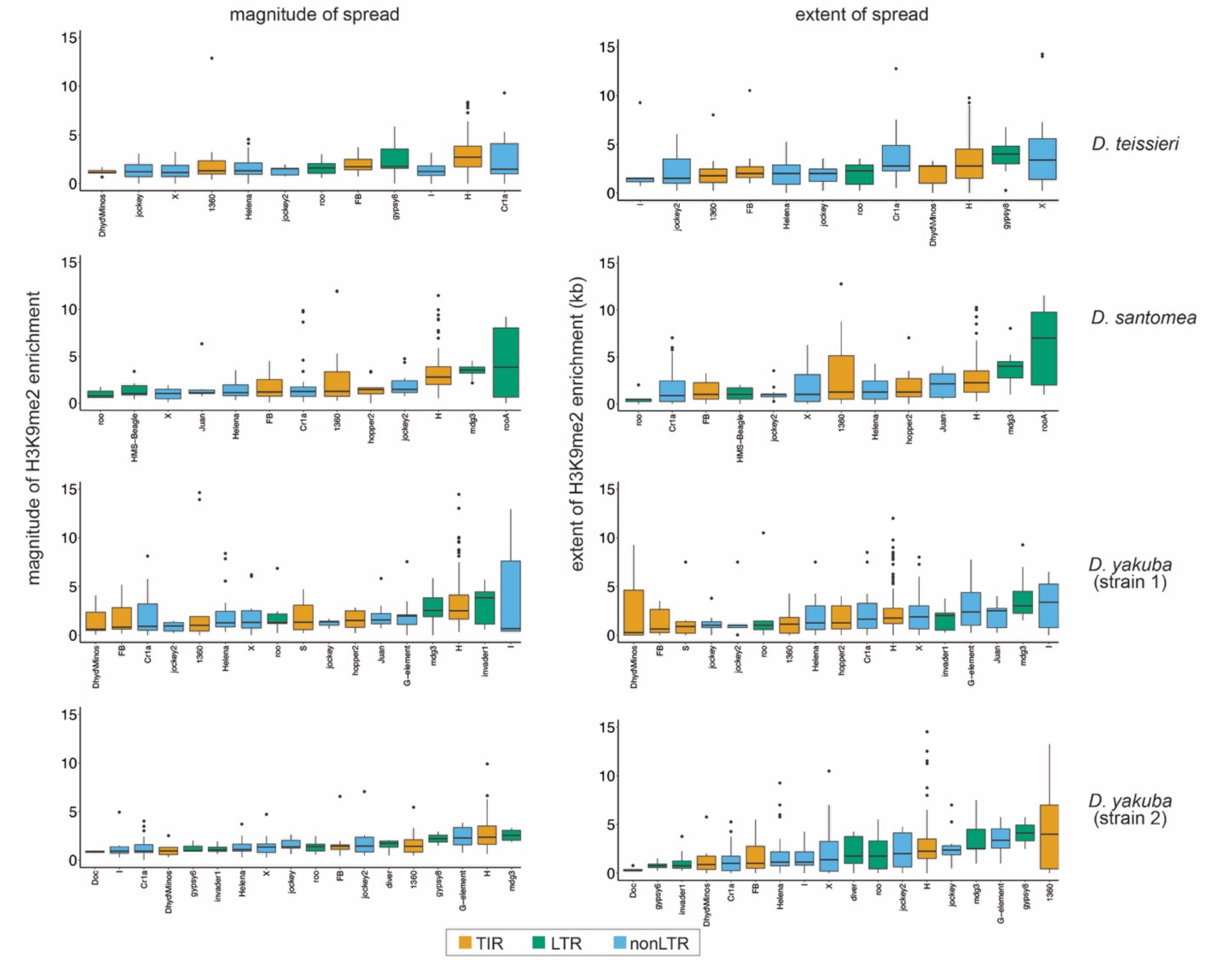

**Figure 2 – Supplementary Figure S3.** The percentage of TEs assigned to a TE family based on blast analysis is shown. This percentage is influenced by the minimum size of TEs (200bp vs 500bp) and is biased against those in species in the *yakuba* complex.

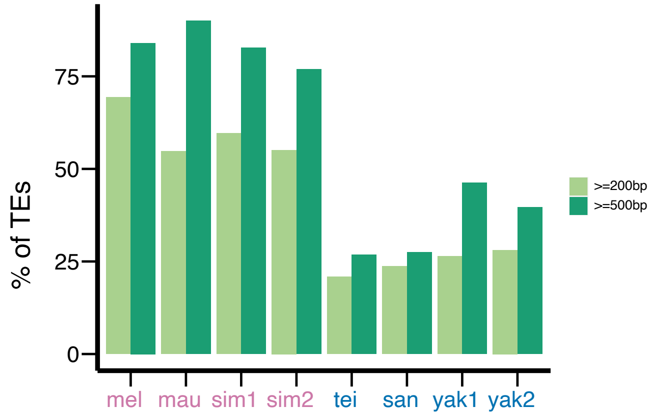

**Figure 3 – Supplementary Figure S1.** *Spearman rank correlation coefficients* ($\rho$) between the magnitude and extent of TE-mediated H3K9me2 enrichment and the H3K9me2 enrichment level of nearby genes. Most of the *Spearman rank correlation coefficients* are significantly different from zero. ***$p < 0.001$, **$p < 0.01$, *$p < 0.05$.

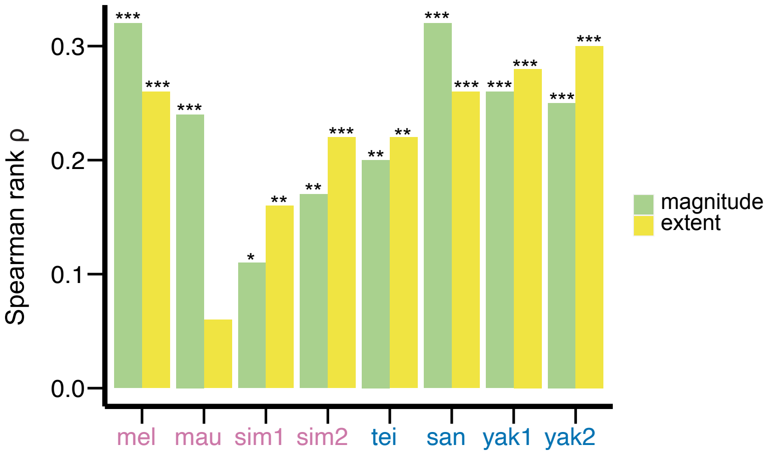

**Figure 3 – Supplementary Figure S2.** The associations between the extent of TE-mediated H3K9me2 enrichment and genic H3K9me2 enrichment differ between genes close (distance to a TE is smaller than the 50% quantile, blue) and distant (gray) to TEs. It is worth noting that, contrary to expectation, the associations are stronger for genes *distant* to TEs than to those close to TEs for *D. melanogaster* and *D. mauritiana*. See Discussions for possible reasons. *Likelihood ratio tests,* ***$p < 0.001$, **$p < 0.01$, *$p < 0.05$.

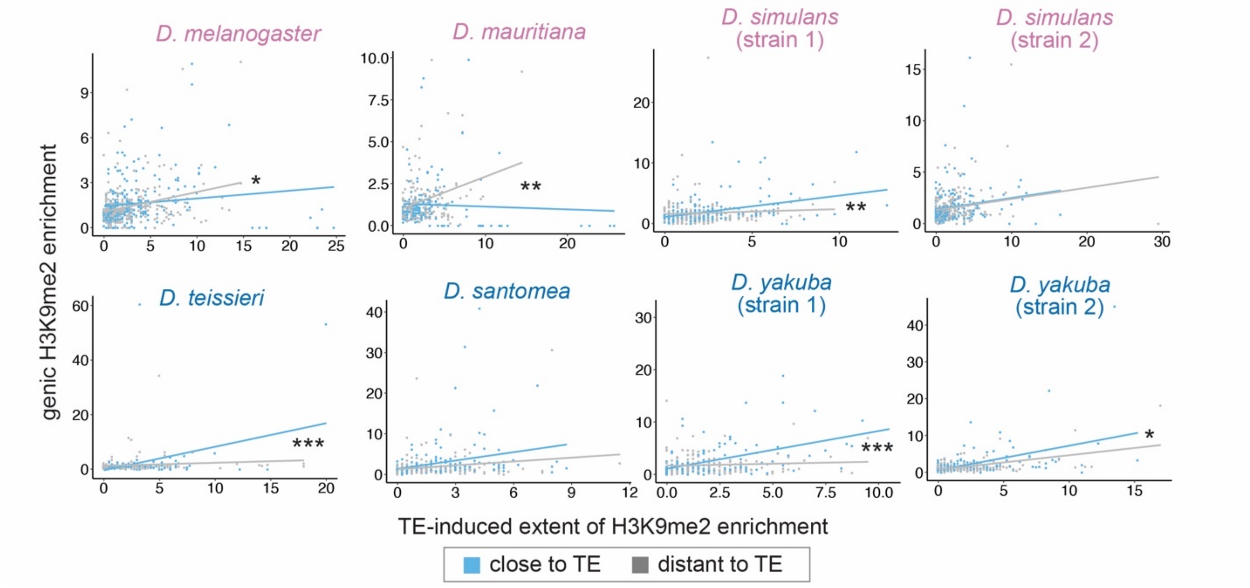

**Figure 3 – Supplementary Figure S3.** The associations between the extent of TE-mediated H3K9me2 enrichment and gene expression rank *does not* differ between genes close (orange) and distant (gray) to TEs. *Likelihood ratio tests* are insignificant ($p > 0.05$) for all genomes.

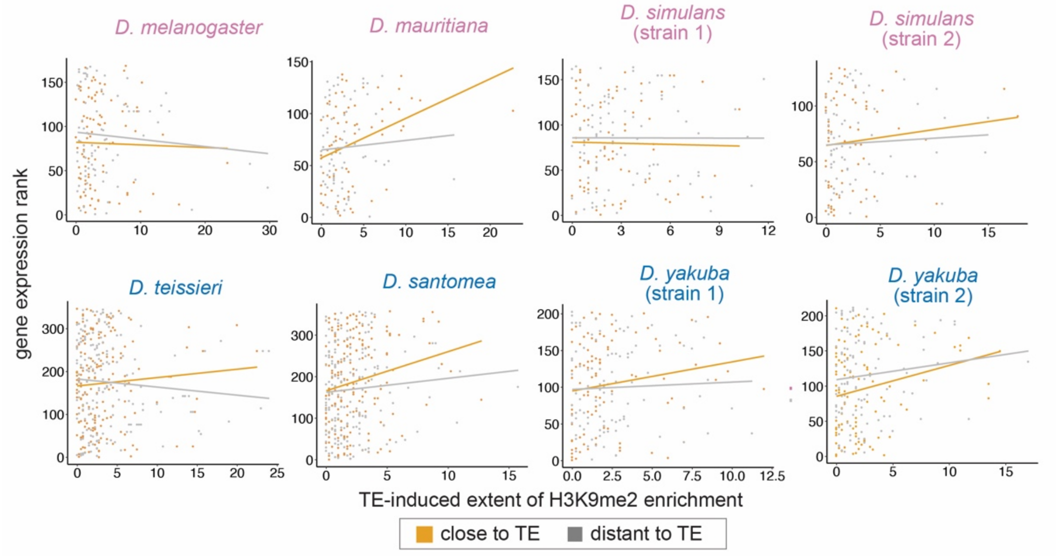

**Figure 3 – Supplementary Figure S4.** Z-scores for comparing the H3K9me2 enrichment (left) and expression rank (right) of homologous genic alleles whose nearby TEs with (blue/orange) or without (gray) epigenetic effects (as defined as the extent of H3K9me2 enrichment > 1kb; see **Figure 3C** for categorizing TEs with the magnitude of H3K9me2 enrichment, which gives similar results). Positive Z-score means the allele with TE has higher H3K9me2 enrichment or larger expression rank (i.e., lower expression level) than the homologous allele without TE in another strain. *Mann-Whitney U test*, ***$p$ < 0.001.

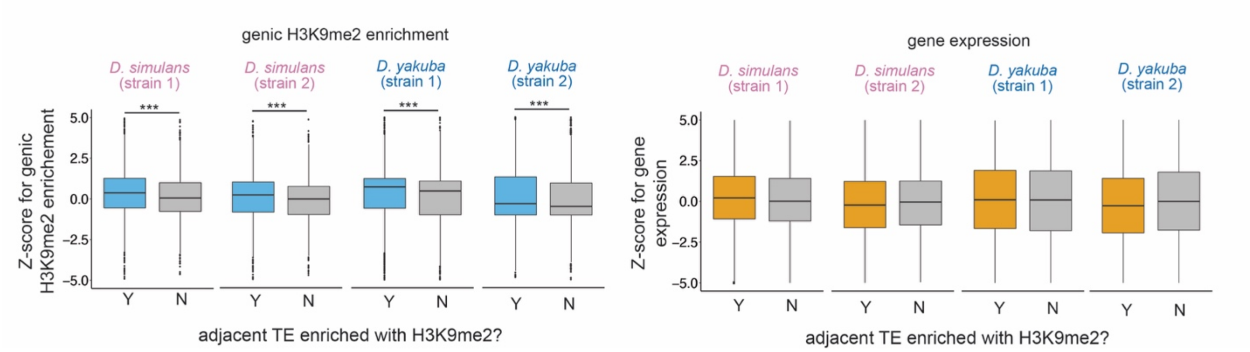

**Figure 3 – Supplementary Figure S5.** Z-scores for comparing the *extent* of TE-mediated H3K9me2 enrichment on the intergenic side and on the genic side for TEs close (green) and far (gray) from genes whose expression is *smaller than* 10 RPKM. The z-scores are not significantly different from zero nor differ between TEs close/distant to genes. *Mann-Whitney tests* are insignificant ($p$ > 0.05) for all comparisons.

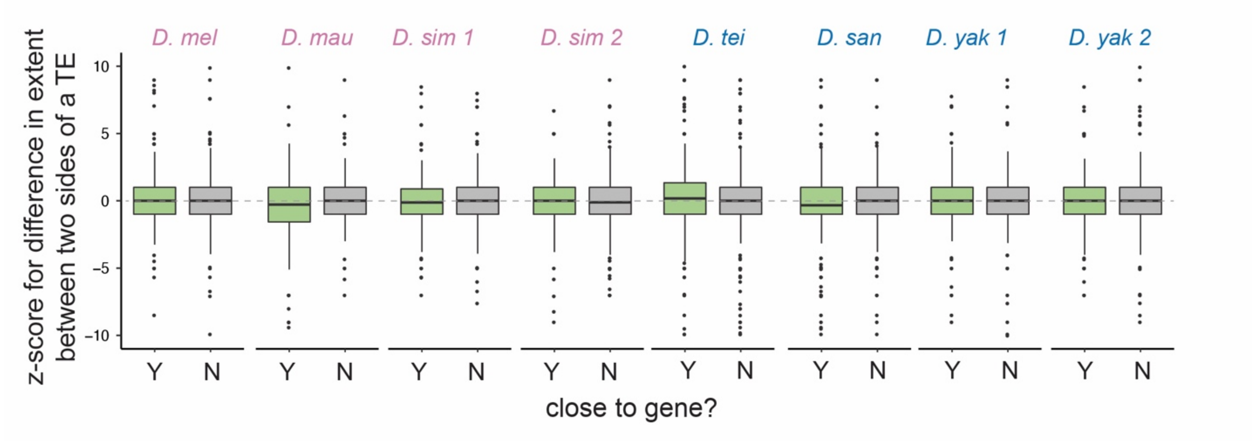

**Figure 3 – Supplementary Figure S6.** Z-scores for comparing the *magnitude* of TE-mediated H3K9me2 enrichment on the intergenic side and on the genic side for TEs close (green) and far (gray) from genes whose expression is at least 10 RPKM. The z-scores are not significantly different from zero nor differ between TEs close/distant to genes. *Mann-Whitney tests* are insignificant ($p > 0.05$) for all comparisons.

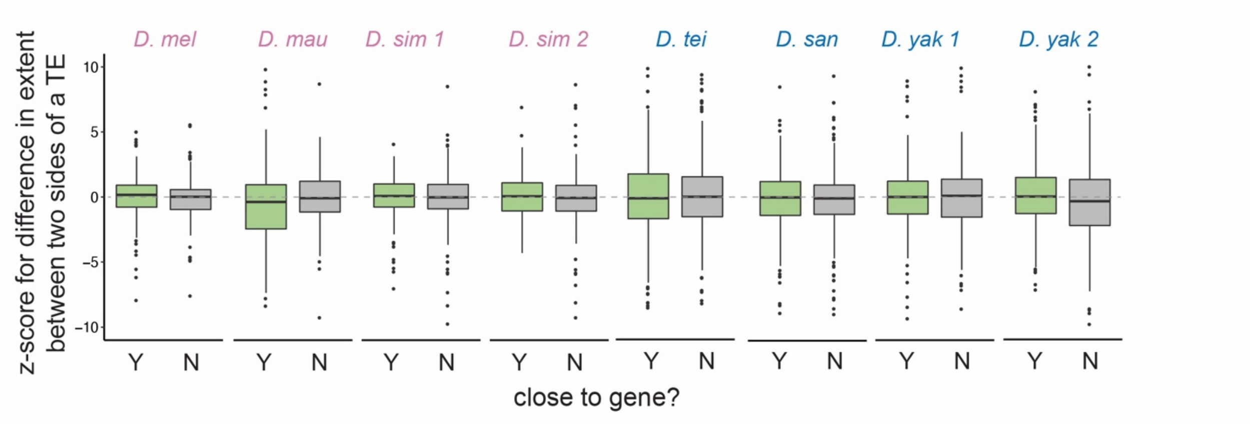

**Figure 3 – Supplementary Figure S7** For TEs near genes, the extent of TE-induced H3K9me2 enrichment is influenced by highly expressed (RPKM > 10), but not by lowly expressed (RPKM < 10) genes. Categorization of genes into high and low expression was based on the expression level in another strain that has different TE insertions to avoid confounding effects. *Mann-Whitney U test*, *$*p$* < 0.05.

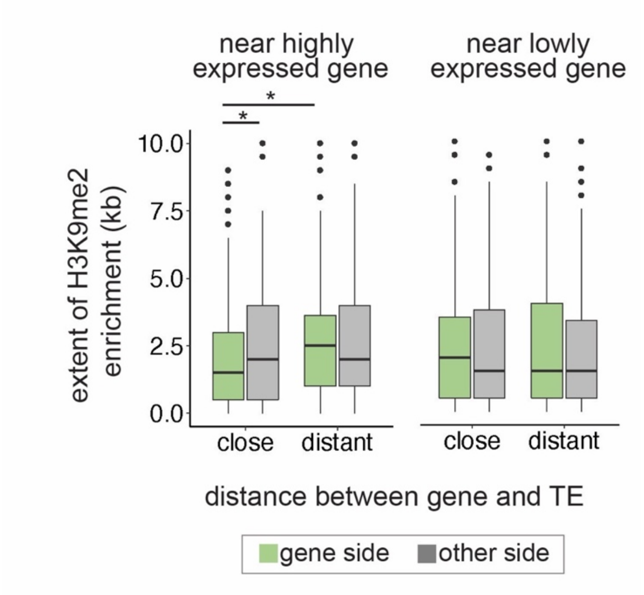

**Figure 3 – Supplementary Figure S8** The extent of TE-induced H3K9me2 enrichment on the side facing insulator sequences CTCF (A) and BEAF-32 (B) or on the other side. In contrast to **Figure 3D,** the extent of H3K9me2 enrichment from TE is similar between sides facing or not facing an insulator sequence, except for one strain of *D. simulans.* CTCF and BEAF-23 sequences are from (Nègre *et al.* 2010). *Paired Mann-Whitney U or Mann-Whitney U test, *p* < 0.05

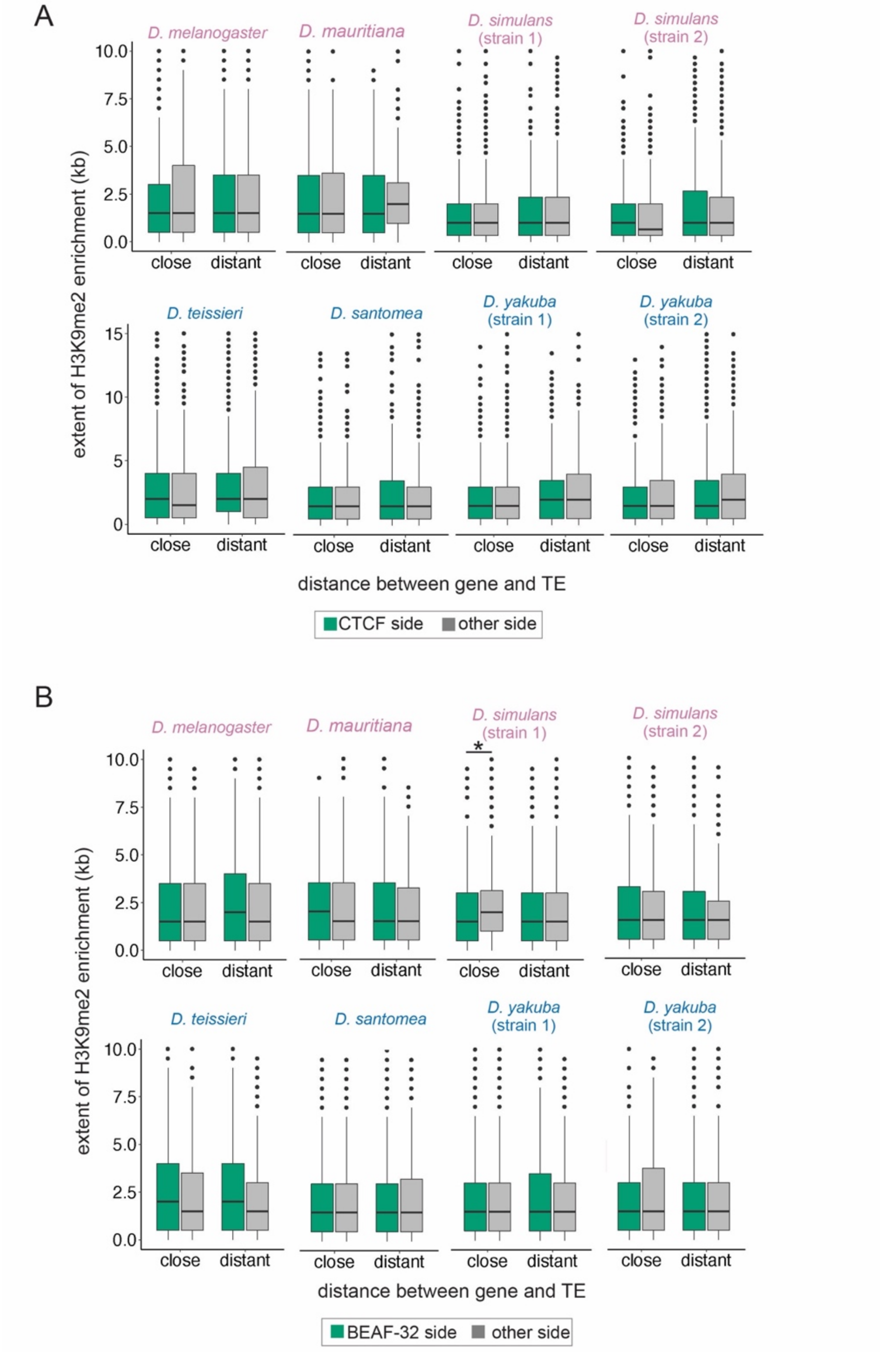

**Table S1:** Correlation coefficient and regression coefficient for the associations between *Su(var)* expression rank and the magnitude of TE-mediated H3K9me2 enrichment. Genome-wide percentiles for correlation coefficient: -0.7142 (5%) and =-0.5952 (10%); genome-wide percentiles for regression coefficient: -0.4208 (5%) and -0.3208 (10%).

| | | magnitude | |
|---|---|---|---|
| **FBgn** | **gene name** | **correlation coefficient** | **regression coefficient** |
| FBgn0000100 | RpLP0 | 0.3333 | -0.3650 |
| FBgn0000662 | fl(2)d | 0.2381 | 0.1767 |
| FBgn0001197 | His2Av | -0.5476 | -0.3186 |
| FBgn0001308 | Khc | -0.1905 | -0.1181 |
| FBgn0002780 | mod | -0.1667 | 0.4934 |
| FBgn0003449 | snf | 0.0000 | -0.2079 |
| FBgn0003598 | Su(var)3-7 | -0.2143 | -0.1797 |
| FBgn0003607 | Su(var)205 | -0.5476 | -0.1203 |
| FBgn0003612 | Su(var)2-10 | 0.2143 | -0.0540 |
| FBgn0004103 | Pp1-87B | -0.4286 | 0.0062 |
| FBgn0004401 | Pep | 0.0238 | -0.1401 |
| FBgn0004655 | wapl | -0.1667 | -0.1581 |
| FBgn0004914 | Hnf4 | -0.1429 | 0.3952 |
| FBgn0005278 | Sam-S | 0.0952 | 0.1033 |
| FBgn0015268 | Nap1 | -0.3571 | -0.2395 |
| FBgn0015396 | jumu | -0.2857 | 0.0406 |
| FBgn0015805 | HDAC1 | -0.2381 | -0.2097 |
| FBgn0020309 | crol | 0.2143 | 0.3173 |
| FBgn0025355 | SuUR | -0.2857 | -0.1956 |
| FBgn0025639 | Hmt4-20 | -0.2857 | 0.1910 |
| FBgn0026427 | Su(var)2-HP2 | -0.5476 | 0.0568 |
| FBgn0026573 | ADD1 | 0.2381 | 0.1094 |
| FBgn0027567 | CG8108 | -0.2381 | -0.2011 |
| FBgn0027835 | Dp1 | -0.5000 | 0.0653 |
| FBgn0027951 | MTA1-like | -0.1667 | -0.2037 |
| FBgn0028387 | chm | 0.1905 | 0.2736 |
| FBgn0030301 | HP5 | -0.5476 | -0.0665 |
| FBgn0033233 | Kdm4A | -0.0952 | -0.1683 |
| FBgn0034217 | Lhr | -0.6905 | 0.2839 |
| FBgn0035829 | HP4 | -0.6190 | -0.1001 |
| FBgn0038551 | Odj | -0.0952 | -0.2191 |

| | | | |
|---|---|---|---|
| FBgn0039338 | XNP | -0.5000 | 0.0612 |
| FBgn0040372 | G9a | 0.0000 | 0.2423 |
| FBgn0085424 | nub | 0.5714 | 0.4850 |
| FBgn0086908 | egg | 0.0714 | -0.2248 |
| FBgn0087035 | AGO2 | 0.7381 | -0.4078 |
| FBgn0260397 | Su(var)3-3 | 0.3810 | -0.4041 |
| FBgn0263144 | bin3 | -0.7381 | -0.4070 |
| FBgn0263755 | Su(var)3-9 | -0.4791 | -0.1569 |
| FBgn0266557 | kis | -0.5476 | -0.1109 |
| FBgn0266599 | Hsc70-4 | -0.6190 | -0.3040 |
